# Histone methyltransferase DOT1L maintains identity and restricts cytotoxic potential of CD8 T cells

**DOI:** 10.1101/2025.01.20.633937

**Authors:** Muddassir Malik, Willem-Jan de Leeuw, Muhammad Assad Aslam, Eliza Mari Kwesi-Maliepaard, Teun van den Brand, Bram van den Broek, Maxime Kempers, Liesbeth Hoekman, Natalie Proost, Tibor van Welsem, Elzo de Wit, Jannie Borst, Heinz Jacobs, Fred van Leeuwen

**Author notes:** shared first author (Muddassir Malik, Willem-Jan de Leeuw). equal contribution (Muhammad Assad Aslam, Eliza Mari Kwesi-Maliepaard).

## Abstract

The histone methyltransferase DOT1L is emerging as a central epigenetic regulator in immune cells. Loss of DOT1L during development of CD8 T cells *in vivo* leads to gain of memory-features but has also been reported to compromise CD8 T cell viability and activity. Here, we determined the cell-intrinsic role of DOT1L in mature mouse CD8 T cells. After conditional deletion of *Dot1L in vitro,* CD8 T cells retained *in vivo* proliferative capacity and anti-tumor reactivity. Moreover, *Dot1L* knock-out CD8 T cells showed increased antigen-specific cytotoxicity towards tumor cells *in vitro.* Mechanistically, loss of DOT1L resulted in an altered cell-identity program with loss of T-cell and gain of NK-cell features. These transcriptional changes were mediated by loss of DOT1L methyltransferase activity in a dose-dependent manner. Our findings show that ablation of DOT1L activity in mature CD8 T cells is well-tolerated and rewires their cell identity towards the NK-cell lineage, concomitantly enhancing intrinsic cytotoxic capacity.

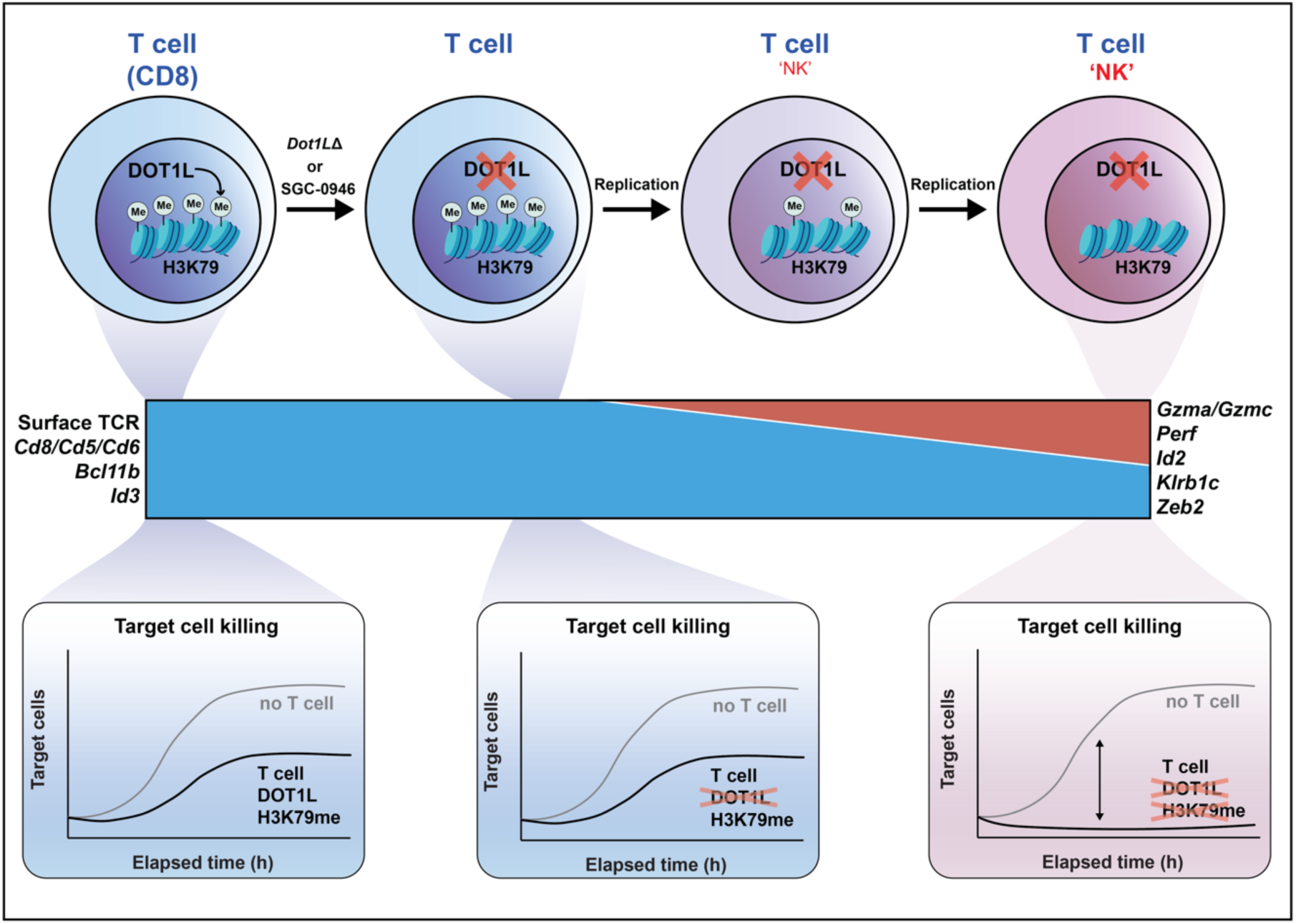

**Highlights:**

- Differentiation and identity of cytotoxic T cells depend on the methyltransferase DOT1L
- DOT1L intrinsically limits the cytotoxic activity of mature CD8 T cells
- DOT1L prohibits the acquisition of NK cell features in CD8 T cells
- DOT1L maintains CD8 T cell identity through its catalytic activity in a dose-dependent manner

## Introduction

Cytotoxic CD8-positive (CD8) T cells are key executors of the adaptive immune system. Their differentiation and activation are tightly regulated by key transcription factors and guided by epigenetic mechanisms (*1–3*). The toolbox for manipulating epigenetic regulators is rapidly expanding, thereby offering new opportunities to influence CD8 T cell fate and function. However, harnessing the regulatory potential of the epigenome to modulate CD8 T cell mediated immunity requires the identification and mechanistic understanding of the key epigenetic regulators involved.

We previously uncovered a central role for the histone methyltransferase DOT1L in mouse CD8 T cell development (*4*). DOT1L methylates histone H3 lysine 79 (H3K79) and ‘writes’ H3K79 methylation (H3K79me) at the 5’end of gene bodies of actively transcribed genes, where the degree of H3K79 dimethylation (H3K79me2) correlates with the level of gene activity (*5*). While the molecular functions of DOT1L and H3K79me are still largely unknown, DOT1L has been recognized as a barrier to differentiation in the immune system (*4, 6–10*). Moreover, highly specific inhibitors have been developed that allow for precision targeting of DOT1L *in vitro* or *in vivo*, as has been demonstrated for tumor cells in which DOT1L activity forms a cell-type specific dependency (*11–14*). However, not all molecular and cellular functions of DOT1L are dependent on H3K79 methylation, indicating that DOT1L might have additional roles beyond its canonical enzymatic function (*15–20*).

Using a conditional mouse knock-out model in which *Dot1L* is deleted early during thymic T-cell development, (Lck-Cre;Dot1L), we observed that *Dot1L* knock-out CD8 T cells prematurely differentiate in an antigen-independent manner towards memory-phenotype cells (referred to as T_AIM_ cells) (*4, 21*). This premature differentiation from naïve T cells towards cells with enhanced memory features suggests that loss of DOT1L may promote more effective and sustained immunological responses (*21–23*). However, in the same *Dot1L* KO mice, the CD8 T-cell response against *Listeria monocytogenes* infection and to vaccination was impaired (*4*). In addition, deletion of *Dot1L* later during thymic T-cell development (CD4-Cre;Dot1L) *in vivo* has been linked to the development of CD8 T cells with molecular changes indicating impaired function (*24*). Mice of this latter model showed reduced control of MC38 tumor cell outgrowth compared to wild-type mice and increased apoptosis in CD8 T cells in tumor draining lymph nodes and tumor tissues (*24*). In these two *Dot1L* knock-out models, *Dot1L* is also deleted in other immune cell types, leading to compromised development of CD4+ T and invariant NKT (iNKT) cells (*4*). Given DOT1L’s impact on various members of the lymphoid lineage, these additional confounding changes may influence the differentiation of CD8 T cells and the overall immune responses in these animals, affecting T cells both directly and indirectly. Finally, treatment of human T cells with DOT1L inhibitor (DOT1Li) SGC-0946 reportedly elevates the activation threshold of CD8 T cells. This elevated threshold reduced the graft versus host response *in vivo* upon transfer of the DOT1Li-treated T cells to immunodeficient NSG mice, while CAR-T cell mediated tumor control activity was maintained (*25*). However, in this chemical perturbation model, DOT1L inactivation is not permanent, with regain of H3K79 methylation and other DOT1L functions expected in absence of DOT1Li following transfer.

To conclude, DOT1L has been ascribed positive as well as negative roles during T-cell development and differentiation. In this study, we combined inducible genetic and chemical conditional inactivation models in mature CD8 T cells to overcome previous confounding factors and thereby understand the cell-intrinsic role of DOT1L in mature CD8 T cells. The results of these models suggest that mature CD8 T cells lacking DOT1L activity are proficient in antigen-specific proliferation and maintain their ability to control tumor growth *in vivo*. Moreover, *Dot1L* KO CD8 T cells showed accelerated and enhanced antigen-dependent tumor cell killing *in vitro*. Mechanistically, this role was mediated by the catalytic activity of DOT1L, as chemical inhibition phenocopied genetic deletion of *Dot1L* in CD8 T cells. Multi-omics phenotyping of the *Dot1L*-KO and DOT1L-inhibited CD8 T cells provided evidence for altered differentiation with partial loss of canonical CD8 T cell features and gain of NK cell properties. These findings suggest that one key cell-intrinsic role of DOT1L is to maintain CD8 T-cell identity and restrict their cytotoxic activity. These insights support further investigations into strategic targeting of DOT1L coupled with other molecular modifications to modulate immune responses *in vivo*.

## Results

### Mature CD8 T cells tolerate loss of DOT1L and H3K79 methylation

In previous studies on DOT1L in CD8 T cells, *Dot1L* was permanently deleted early in the T cell lineage or transiently inactivated by chemical inhibition (*4, 24–26*). To overcome the limitations of those models and to determine the cell-intrinsic role of DOT1L in mature CD8 T cells, we generated a tamoxifen (4-OHT)-inducible deletion model (Cre-ERT2;Dot1L^fl/fl^) allowing genetic ablation of *Dot1L* in spleen-derived CD8 T cells *in vitro* (Fig. 1A). In this model, CD8 T cells also expressed the transgenic OVA-specific T-cell receptor (TCR) OT-I (*27*), which recognizes the chicken ovalbumin (OVA) derived SIINFEKL peptide in the context of H-2K^b^.

**Figure 1.**
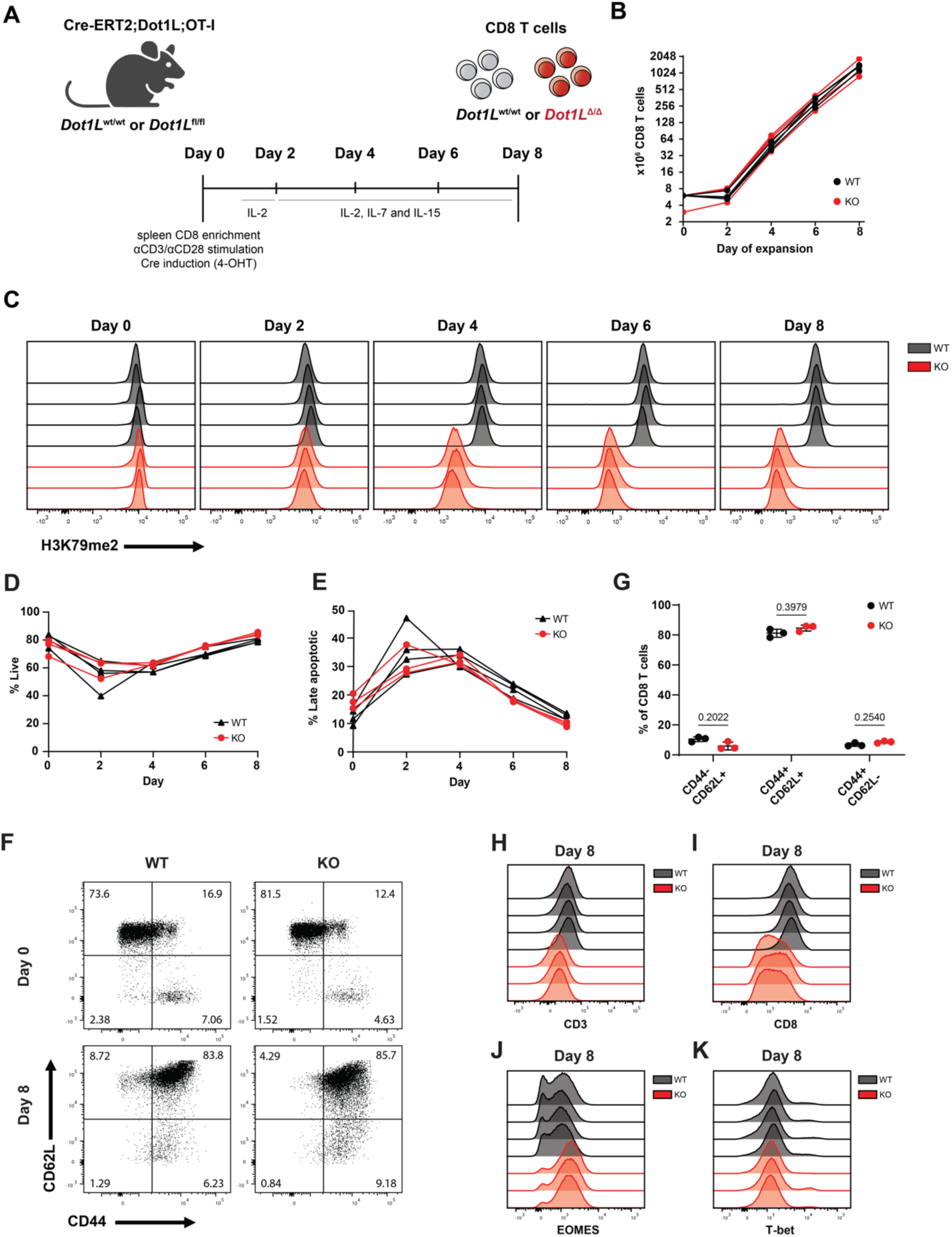
*Dot1L* deletion in mature CD8 T cells leads to replication-dependent loss of H3K79me2 without compromising viability. (A) Experimental set-up: Mature CD8 T cells were harvested from spleens of mice expressing tamoxifen-inducible Cre recombinase (CreERT2) and the OT-I transgenic TCR and carrying either wild-type (wt) or floxed (fl) *Dot1L* alleles. CD8 T cells were activated (anti-CD3/CD28), treated with 4-OHT to induce Cre, and expanded in the presence of cytokines as indicated. (B) Expansion of *Dot1L*^wt/wt^ (WT) and *Dot1L*^Δ/Δ^ (KO) CD8 T cells. (C) Flow cytometry analysis of KO-induced loss of H3K79me2 over time; histograms with n=4 (WT) and 3 (KO) biological replicates. (D-E) Flow cytometry analysis of viability (D) and late apoptosis (E); CD8 T cells were gated on AnnV-/DAPI-for live and AnnV^-^/DAPI^+^ for late apoptotic/necrotic cells (WT n=4 and KO n=3 biological replicates). (F) Flow cytometry analysis dot plots of differentiation status of WT and KO CD8 T cells at Day 0 and Day 8 based on CD44 and CD62L expression; representative biological replicates plotted. (G) Quantification of CD8 T cell subsets as in panel F (average of n=3 biological replicates +/− SD; 2-way ANOVA and Šídák’s multiple comparisons test). (H-K) Histograms of surface expression of CD3, CD8, and nuclear staining for T-bet and EOMES based on flow cytometry.

Isolated splenic OT-I CD8 T cells were activated with CD3 and CD28 agonists, treated with 4-OHT and expanded for eight days *in vitro* (Fig. 1A-B). CD8 T cells derived from Cre-ERT2;Dot1L^wt/wt^ mice were used as wild-type controls (WT). While genomic deletion of exon 2 of *Dot1L* was efficient and nearly complete at Day 2 (Supplementary Fig. S1A), H3K79me was gradually lost over time, as determined by flow cytometry analysis of H3K79me2 (Fig. 1C). The slow decay of the H3K79me mark agrees with the absence of a known H3K79-specific demethylase. Loss of H3K79-methylation is therefore mainly determined by passive dilution upon cell proliferation (*28, 29*). H3K79me2 staining was strongly reduced after four days of T cell expansion and stabilized around the lowest level on Day 6 (Fig. 1C and Supplementary Fig. S1B). Mass spectrometry analysis further confirmed that in *Dot1L* KO cells on Day 8, no methylation was detectable, indicating that DOT1L is the sole methyltransferase for H3K79 in T cells (Supplementary Fig. S1C and Kwesi-Maliepaard *et al.* (*4*)).

Despite the rapid deletion of *Dot1L* and subsequent passive loss of H3K79me, *Dot1L* KO CD8 T cells expanded at a similar rate as WT cells (Fig. 1B). Moreover, no loss of viability was observed (Fig. 1D-E), consistent with previous results (Kwesi-Maliepaard et al. 2020) but contrasting observations when *Dot1L* is deleted early in the T cell lineage (*24*). We next analyzed the expression of key CD8 T cell surface proteins and transcription factors. In our experimental set-up, naïve cells acquired CD44 expression following activation. Deletion of *Dot1L* did not affect this differentiation pattern (Fig. 1F-G). Surface levels of CD3 (TCR complex) and the CD8 co-receptor were reduced upon *Dot1L* knockout in mature T cells, potentially indicating partial loss of key CD8 T cell specific markers (Fig. 1H-I and Supplementary Fig. S1D). Finally, loss of DOT1L led to increased expression of the CD8 T cell transcription factor EOMES, while T-bet was unaffected (Fig. 1J-K, Supplementary Fig. S1E-G). These observations partially recapitulated the characteristics of *Dot1L* deficient CD8 T_AIM_ cells in the Lck-Cre;Dot1L model, in which reduced CD8 and CD3 surface expression and increased EOMES and T-bet expression were observed (*4*). T-bet and EOMES are two transcription factors that determine effector CD8 T cell differentiation and function (*3, 30*). These findings show that, apparently, loss of DOT1L modulates key aspects CD8 T cell identity without impairing their viability.

### Proficient tumor control *in vivo* by *Dot1L* KO CD8 T cells upon adoptive cell transfer

Mice in which *Dot1L* is deleted early in the T cell lineage show reduced immune responses (*4, 24*), including compromised tumor control (*24*). In line with this observation, impaired CD8 T cell functionality due to limited methionine availability in the tumor microenvironment has been linked to reduced DOT1L activity (*24*). To address whether the observed loss of functionality *in vivo* upon early deletion of *Dot1L* reflects a cell-intrinsic role of DOT1L, we here used an in *vitro* deletion approach in mature CD8 T cells. To determine the anti-tumor potential of mature CD8 T cells lacking DOT1L, B16F10 melanoma cells expressing OVA (B16F10-OVA) were subcutaneously injected in CD45.1+ recipient mice to initiate tumor outgrowth. After eight days of tumor outgrowth, the tumor-bearing mice received adoptive cell therapy (ACT) with *in vitro* expanded, 4-OHT-treated CD45.2+ OT-I CD8 T cells Fig. 2A). Loss of H3K79me2 as well as reduced surface expression of the TCR and CD8 co-receptor on Day 8 of *in vitro* expansion of *Dot1L* KO T cells was confirmed by flow cytometry (Fig. 2B-D and Supplementary Fig. S2A-B). Mice treated with vehicle (HBSS) had a median survival of 27 days, while mice that received WT CD8 T cells had a significantly increased median survival of 42 days (Fig. 2E). However, no difference in survival was observed between WT and KO ACT treatment (median survival of 44 days; (Fig. 2E). Similarly, the cumulative tumor outgrowth was significantly delayed after ACT of WT (mean: 38.4±4.9 days) as well as KO (mean: 39.1±5.8 days) CD8 T cells compared to vehicle control (mean: 25.0±7.3 days) but was not different between KO and WT (Fig. 2F; Supplementary Fig. S2C). No significant endpoint bias could be observed based on tumor weight at the time of sacrifice (Supplementary Fig. S2D).

**Figure 2.**
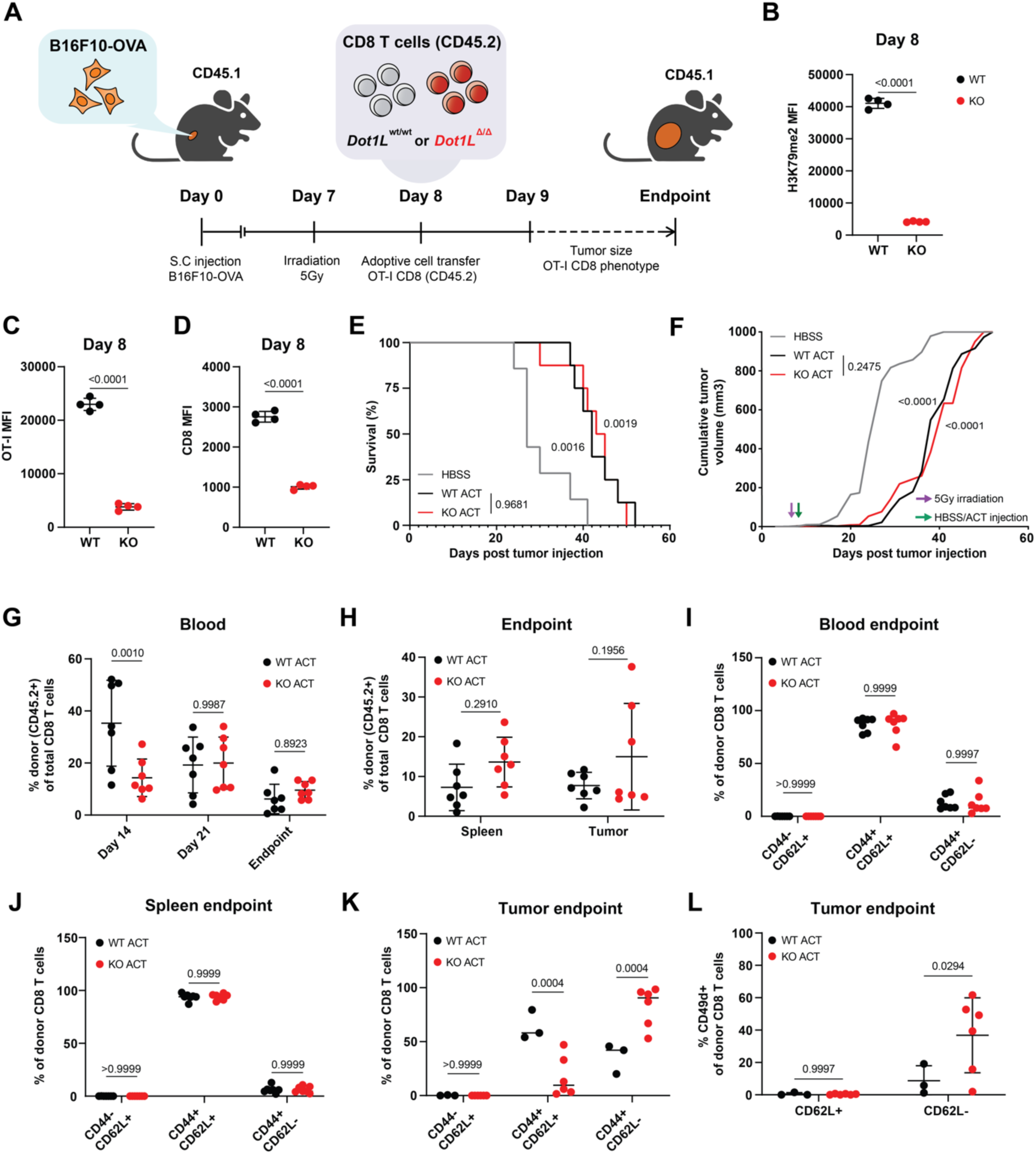
Adoptively transferred *Dot1L* KO CD8 TCR-T cells reduce tumor outgrowth *in vivo*. (A) Experimental setup: At Day 0, CD45.1+ recipient animals were subcutaneously injected with B16F10 melanoma cells expressing OVA. Mice were irradiated (5Gy) at Day 7 and received HBSS (vehicle), CD45.2+ WT, or KO OT-I CD8 T cells at Day 8. CD45.2+ OT-I CD8 T cells were expanded as in Fig. 1A. (B-D) Flow cytometry quantification of H3K79me2 (B) levels, and OT-I (C) and CD8 surface expression (D) of expanded donor CD8 T cells, prior to adoptive cell transfer (average of n=4 biological replicates +/− SD; unpaired t-test). (E) Survival plot of tumor-bearing mice following ACT. Animals were sacrificed at endpoint, which was based either on welfare or tumor size (>1000 mm^3^). Significance was determined by a Log-rank (Mantel-Cox) tests between groups (n=7 (HBSS) and n=8 (WT and KO)), Bonferroni-corrected α value= 0.05/3, thus significant if P<0.0167). The survival logrank hazard ratio (HR) for both WT (HR: 0.2692, 95%CI: 0.07423-0.9763) and KO (HR: 0.2775, 95%CI: 0.07744-0.9945) was lower compared to vehicle treated mice, but not different between KO ACT to WT ACT treated mice (HR: 1.023, 95%CI: 0.3821-3.057). (F) Cumulative tumor growth over time after ACT (n=7). Significance was determined by cumulative Gaussian curve fitting and best-fit analyses. (G) Percentage donor CD8 T cells of total CD8 T cells in blood at indicated time points for WT and KO ACT mice (flow cytometry, average of n=7 +/− SD). (H) Percentage donor CD8 T cells in spleen and tumors at endpoint (average of n=7 +/− SD). Blood (see G) at endpoint was included in statistical analysis but not displayed. (I-K) Differentiation state of T cells at endpoint based on CD44 and CD62L staining in (I) blood and (J) spleen (average of n=7 +/− SD). (K) Differentiation state of donor CD8 T cells in tumor at endpoint based on CD44 and CD62L flow cytometry staining. (L) Percentage CD49d+ donor CD8 T cells in tumor at endpoint (flow cytometry), grouped by CD62L expression (n=3 (WT) and n=6 (KO), average +/− SD; tumor samples with less than 150 events in the CD45.2+ lymphocyte gate were excluded). For G-L, significance was determined by paired 2-way ANOVA and Šídák’s multiple comparisons tests.

In addition to monitoring tumor growth, we analyzed the abundance and phenotype of transferred CD45.2+ OT-I CD8 T cells isolated from blood (Day 14 and 21 and endpoint), and the spleen and tumor tissue at endpoint (time of sacrifice, which was variable between individual animals). Although at Day 14, the relative abundance of KO cells in blood was lower than that of WT transferred cells, WT and KO donor CD8 T cells expanded to the same relative abundance at Day 21 in blood, and at end point in spleen and tumor tissue (Fig. 2G-H). Regarding the phenotype, the immune composition in blood at spleen at end point was predominantly made up of CD44 and CD62L (L-selectin) double-positive cells with no significant differences observed between WT and KO (Fig. 2I-J). In contrast, in the tumor at endpoint, the *Dot1L* KO tumor-infiltrating lymphocytes (TILs) showed a gain of the CD44+CD62L- and loss of the CD44+CD62L+ population compared to WT, indicating increased differentiation towards effector-like cells (Fig. 2K). Moreover, their expression of CD49d, a marker of cognate antigen experience, was higher, as was the percentage of CD49d+ TILs (Fig. 2L and Supplementary Fig. S2F). It should be noted, however that the number of CD45.2+ TILs at endpoint was generally low (Supplementary Fig. S2E), which likely relates to the rapid outgrowth of this aggressive tumor model.

Together, these results show that expansion, viability, and *in vivo* tumor control by adoptively transferred CD8 T cells were not negatively affected by deletion of *Dot1L* and concomitant loss of H3K79me (Figs. 1-2). Moreover, at the endpoint in the B16F10-OVA tumors, *Dot1L* KO cells adapted a more effector-memory like state and increased upregulation of CD49d. To investigate whether these changes reflect a possible role of DOT1L in restricting activation and effector functions of mature CD8 T cells, we focused our further analyses on studying their cytotoxic function *in vitro*.

### Deletion of *Dot1L* increases the tumor-killing potential of mature CD8 T cells *in vitro*

To quantitatively measure their cytotoxic potential, *in vitro* expanded *Dot1L* WT and KO OT-I CD8 T cells (Day 8) were co-cultured with mCherry expressing B16F10-OVA cells (Fig. 3A). Tumor cell killing was determined by quantifying the confluency of mCherry+ B16F10-OVA target cells using 4h interval imaging over a time-course of four days. Under these conditions, wild-type OT-I CD8 T cells showed dose-dependent killing of target tumor cells over time. In comparison, *Dot1L* KO T cells showed enhanced control of target cell outgrowth (Fig. 3B-C). Moreover, at the highest CD8 T-cell to target-cell ratio, the killing was most pronounced, as target cell clearance was near complete within the first 8h of co-culture (Fig. 3B-C, Supplementary Fig. S3A-C, Supplementary Mov. S1-S3).

**Figure 3.**
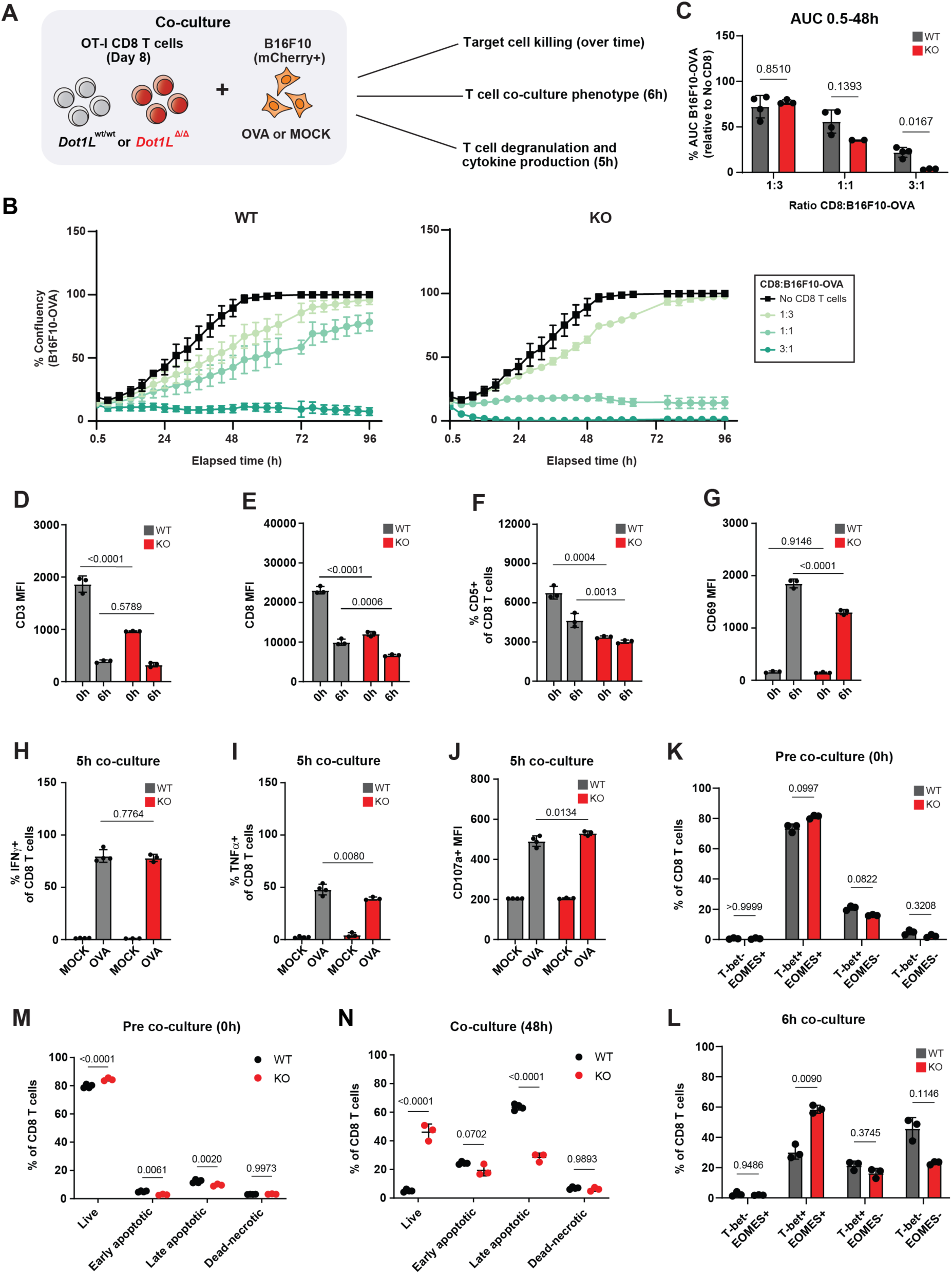
Deletion of *Dot1L* increases the cytotoxic activity of CD8 T cells towards tumor cells *in vitro*. A) Experimental set-up: expanded WT and KO CD8 T cells were co cultured with B16F10-OVA or B16F10-MOCK target cells to determine target cell killing and CD8 phenotypes. (B) Quantification of live cell imaging of mCherry+ B16F10-OVA tumor cell outgrowth during co-culture with different ratios of OT-I CD8 T cells to target cells (B16F10-OVA) or no CD8 cells. Average of biological replicates (n=3 (KO) or n=4 (WT) of expanded CD8 T cells +/− SD). The no-CD8 curves displayed in KO and WT are identical. (C) Quantification of area under the curve (AUC) of (B), relative to no CD8 (AUC at 48h; average +/− SD of biological replicates, n=4 (WT) or n=3 (KO)). (D-G) Quantification of flow cytometry surface expression of CD3 (D) and CD8 (E), %CD5+ (F) and surface expression of CD69 (G) of CD8 T cells prior to co-culture (0h) and after 6h co-culture with B16F10-OVA cells. (H-J) Flow cytometry-based quantification of %IFNγ+ (H), %TNFα+ (I) and CD107a (J) of CD8 T cells after co-culture with OVA or MOCK B16F10 cells for 5 hours, average +/− SD (n=3). (K-L) Quantification of T-bet and EOMES populations by flow cytometry prior to co-culture (K) and after 6h co-culture (I), average +/− SD (n=3). Significance was determined by two-way ANOVA and Šídák’s multiple comparisons test. (M-N) Quantification of apoptotic state of OT-I CD8 T cells by flow cytometry, prior to co-culture (M) and after 48h with B16F10-OVA (N), average +/− SD (n=3 (KO) and n=4 (WT) biological replicates). For M-N, significance was determined by matched two-way ANOVA and Šídák’s multiple comparisons tests.

To gain further phenotypic insight into the enhanced killing by *Dot1L* KO CD8 T cells, WT and KO cells were analyzed just prior to (0h) and during (6h) co-culture by flow cytometry. After exposure to target cells, WT T cells downregulated their TCR (CD3; a TCR-complex member) and co-receptor (CD8) surface expression (Fig. 3D-E), as expected for CD8 cells that are exposed to their cognate antigen (*31*). KO T cells showed lower CD3 and CD8 surface expression at 0h, as observed earlier, and this further decreased at 6h to levels comparable to (CD3) or lower (CD8) than WT (Fig. 3D-E). The surface expression of CD5 was also reduced in KO CD8 T cells, as was the CD5+ fraction before and during co-culture (Fig. 3F, Supplementary Fig. S3D). Since CD5 is a negative regulator of T cell receptor signaling that has been suggested to be part of a feedback loop (*32*), the lower expression of CD5 might reflect a compensatory mechanism for the reduced TCR and CD8 surface levels. After 6h of co-culture, the upregulation of CD69, an activation marker and metabolic gatekeeper of CD8 T cells (*33*), was reduced in *Dot1L* KO compared to WT (Fig. 3G), although the fraction of CD8 T cells that express CD69 was similar (Supplementary Fig. S3E). This suggests that *Dot1L*-KO CD8 T cells show reduced activation strength.

Effector cytokine release and cytolytic markers were measured by flow cytometry to provide insight into the increased effector function in CD8 T cells during the response against target cells. Production of IFNγ was not altered (Fig. 3H, Supplementary Fig. S3F,I), while the production of TNFα was reduced for KO CD8 T cells (Fig. 3I, Supplementary Fig. S3G,J), the latter of which has also been observed for CD8 T cells in the CD4-Cre;Dot1L model (*24*). Moreover, CD107a (LAMP-1), a marker for active degranulation (*34*), was increased at in KO after 5h of co-culture (Fig. 3J), without affecting the fraction of cells that was actively degranulating (CD107a^+^) (Supplementary Fig. S3H,K), indicating increased granulation potential in *Dot1L* KO CD8 T cells. However, the abundance of cytolytic enzyme Granzyme B (GzmB) was similar between WT and KO CD8 T cells (Supplementary Fig. S3L). We confirmed that the activation of OT-I CD8 T cells upon challenge with target cells was strictly antigen dependent, since no phenotypic changes were observed when T cells were co-cultured with B16F10 cells lacking OVA expression (MOCK) (Fig. 3H-J, Supplementary Fig. S3I-L) and no killing of these MOCK target cells was observed (see below).

To explore the underlying molecular mechanism of enhanced killing response of KO T cells, two key transcription factors, T-bet and EOMES were analyzed. Upon challenge with target cells, WT CD8 T cells shifted from a mainly T-bet EOMES double positive state to a double negative or T-bet single positive state. This response was altered in *Dot1L* KO, in which T cells tended to retain the double positive state (Fig. 3K-L). The impact on differentiation upon target cell challenge was further determined by staining for CD44 and CD62L. From 0h to 6h, WT CD8 T cells shifted from mainly (CD44^int^CD62L^+^) to more-intermediate and terminal effector like states (CD44^high^CD62L^int^ and CD44^int^CD62L^-^, respectively). In *Dot1L* KO T cells, this differentiation shift was skewed towards the intermediate differentiation state (Supplementary Fig. S3M-P).

Finally, we observed that the viability of the CD8 T cells (percentage of live cells) was comparable between WT and KO prior to co-culture (Fig. 3M). However, significantly more viable *Dot1L* KO than WT CD8 T cells were present after 48h of exposure to B16F10-OVA tumor cells (Fig. 3N and Supplementary Fig. S3Q). This increase in cell number might be caused by increased survival of activated *Dot1L* KO T cells, or because the accelerated elimination of target cells led to reduced activation-induced cell death. Combined, these results indicate that DOT1L-deficient mature CD8 T cells acquire a molecular state with altered T cell activation properties and differential expression of key CD8 T cell surface proteins, coinciding with enhanced cell-intrinsic cytotoxic killing potential and increased survival.

### CD8 T cells lacking DOT1L acquire NK-like features and upregulate cytolytic factors

To gain a deeper and unbiased understanding of the altered state of *Dot1L* KO CD8 T cells, we determined the transcriptomic changes upon *Dot1L* deletion in mature CD8 T cells after 8 days of culture. RNA sequencing analysis revealed that 1289 genes were upregulated and 1309 downregulated upon *Dot1L* deletion (Fig. 4A). When subjected to Gene Set Enrichment Analysis (GSEA) based on Gene Ontology (GO), KO CD8 T cells showed a negative enrichment for genes associated with ‘T cell receptor signaling’ (Fig. 4B). This potentially agrees with the reduced surface expression of TCR and CD8 observed by flow cytometry. A positive enrichment was found for genes associated with ‘immune response to tumor cells’ and ‘leukocyte degranulation’ (Supplementary Fig. S4A), in agreement with the increased killing potential and increased levels of degranulation marker CD107a during the response against target cells *in vitro*. Furthermore, based on cell-type specific curated gene sets, GSEA analysis showed a positive enrichment for genes annotated as ‘antigen response’ in KO CD8 T cells (*35*), despite lack of antigen exposure (Fig. 4C). Apparently, *Dot1L*-KO CD8 T cells acquire a more effector-memory like state, potentially contributing to the observed rapid killing *in vitro*. Strikingly, KO CD8 T cells were enriched for gene signatures related to bona fide natural killer cells (*36*) and CD8 T cells that lose their T cell lineage commitment and differentiate towards NK-like T cells (T to NK CD8 T cells) (*37*) (Fig. 4C). Combined, these findings suggest that mature T cells lacking DOT1L downregulate some of the key T-cell features (lower TCR, CD8; reduced activation (based on CD69)), and partially gain properties of NK cells (elevated CD107a, increased T to NK and NK-cell signature genes), that could contribute to a gain in effector potential.

**Figure 4.**
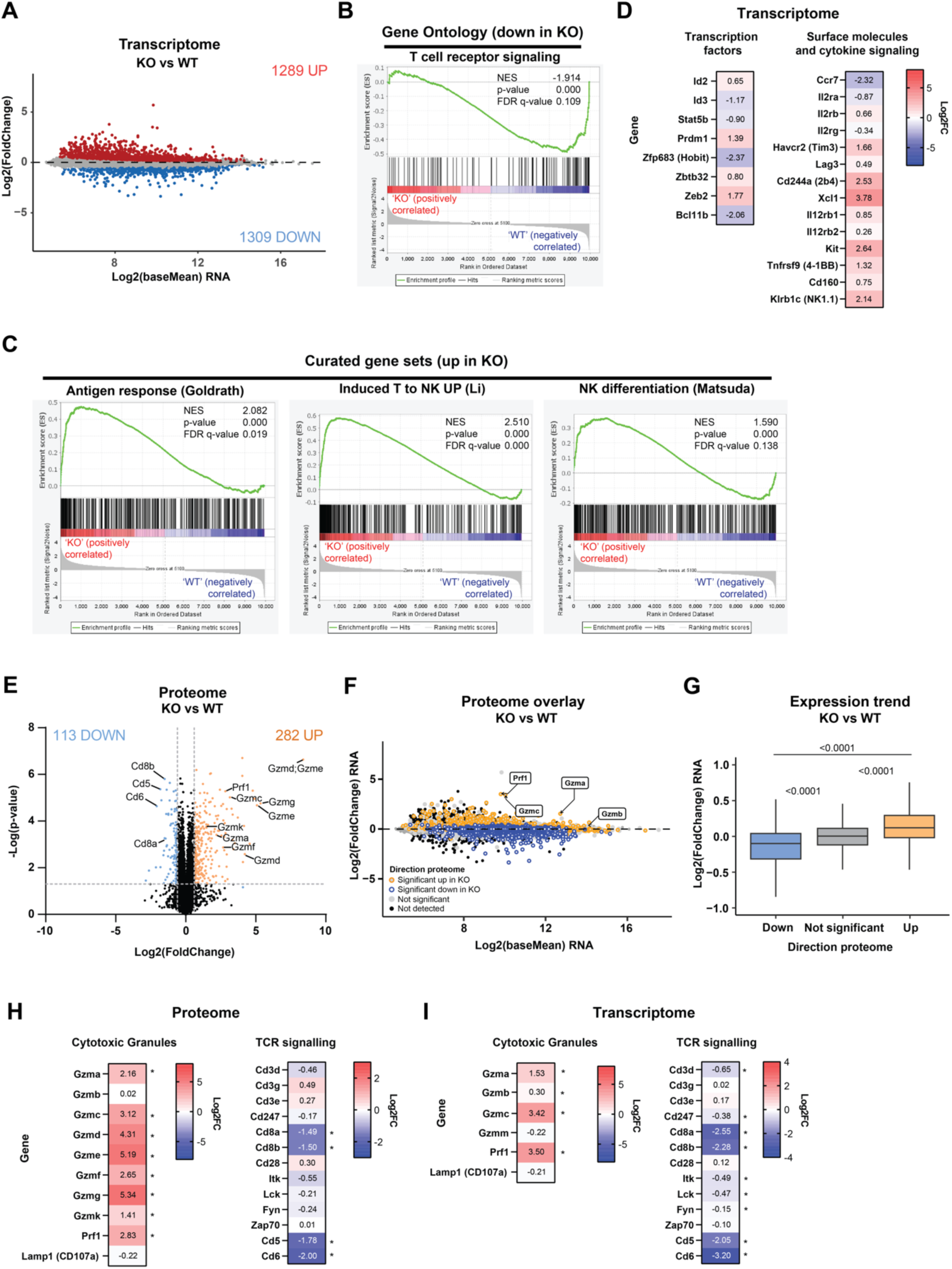
Loss of DOT1L leads to upregulation of key effector proteins in CD8 T cells. (A) Transcriptome analysis of expanded WT and KO CD8 T cells. MA plot shows differentially expressed genes between KO and WT CD8 T cells on Day 8 (average of 3 biological replicates). (B) Gene set enrichment analysis (GSEA) based GO enrichment for T cell receptor signaling. (C) GSEA-based curated enriched gene sets. In (B) and (C), normalized enrichment scores (NES), p-values (Padj) and false discovery (FDR) values are displayed. (D) Transcriptome heatmaps displaying log2 fold-change in KO vs WT for selection of significantly altered (Padj) surface and cytokine signaling molecules and transcription factors (n=3). Positive log2FC refers to an increase in KO. (E) Proteomics volcano plot of differentially expressed proteins between KO and WT CD8 T cells at Day 8 (n=4 (KO) and n=3 (WT) biological replicates), dotted lines represent cut-off values. Highlighted are upregulated cytotoxic granule and downregulated TCR signaling proteins. (F) Overlay of proteome on MA plot of transcriptome. Upregulated (yellow) and downregulated (blue) protein expression for KO CD8 T cells are colored, *Prf1*, *Gzma*, *Gzmb* and *Gzmc* are highlighted. (G) Box plot showing the mRNA changes of proteins that were either differentially downregulated (blue), upregulated (yellow), or not significantly changed (grey) in KO vs WT. Box plot showing the transcriptional trend of three sets of genes that were differentially downregulated or upregulated in KO compared to WT or were non-significant in proteome analysis. Wilcoxon pair wise statistical analysis p-values are plotted. (H-I) Heatmaps of selected proteins (H) and genes (I) involved in cytotoxic granules and TCR signaling (n=3, biological replicates). Genes that were not detected or excluded in the analysis (based on criteria described) were left out of plot. Asterisk represents adjusted P-value (Padj) significance.

These molecular findings guided us to focus on known regulators of T and NK cell differentiation to provide more insight into the altered state of *Dot1L* KO CD8 T cells (Fig. 4D). *Dot1L* KO CD8 T cells show increased expression of *Id2* and *Prdm1* and reduced expression of *Id3*. ID2 (encoded by *Id2*) and ID3 (encoded by *Id3*) are transcription factors known to play a complex co-dependent role in CD8 T cell differentiation and are essential in the formation of effector and memory subsets (*3*). Id3 is essential for memory CD8 T cell formation, and its expression of Id3 is transcriptionally repressed by Blimp-1 (encoded by *Prdm1*) in effector CD8 T cells (*3, 38*). Blimp-1 and Hobit (Homolog of Blimp-1 in T cells, encoded by *Zfp683*) are expressed throughout effector and memory cells and upon antigen stimulation Hobit is downregulated in human CD8 T cells (*39*). Notably, Hobit expression was also downregulated in KO. *Zeb2* was increased in *Dot1L* KO CD8 T cells. ZEB2 (encoded by *Zeb2*) is known to be essential in the CD8 T cell response and known to be a positive regulator in memory and effector memory differentiation (*40*). CCR7 (encoded by *Ccr7*), an important chemokine receptor for T cell homing and typically highly expressed in naïve CD8 T cells (*41, 42*), was downregulated in KO CD8 T cells (Fig. 4D). Together, these changes suggest that loss of *Dot1L* induces several transcriptional programs associated with activated and effector memory CD8 T cells.

Considering the cytotoxic capacity, CD160 (encoded by *Cd160*) expression is associated with cytolytic activity and active effector responses of CD8 T cells; it is also linked to upregulation of 4-1BB (encoded by *tnfrsf9*) (*43, 44*). *Dot1L*-deficient CD8 T cells, which displayed enhanced effector function, showed increased expression of both *Cd160* and *Tnfrsf9* (Fig. 4D).

Interestingly, several of the upregulated factors in KO (e.g. *Id2*, *Zeb2*, *Il2rb*, *Ccr7*, *Tim3*, *Lag3*, *2b4* and *Xcl1*) also play important roles in NK cell differentiation, function and fitness (*40, 45–48*). Furthermore, in the context of NK differentiation, *Il2rb* (CD122), *Cd244* (2b4) and *Klrb1c* (encoding the NK receptor NK1.1 (in humans CD161), all upregulated in KO, are markers often associated with mature NK cells (*45*), and are found on several mouse NK precursors (*49*). For conventional mature NK cells, the expression of Blimp-1 is high and Hobit is low (*39*), matching the observations in KO. Lastly, *Bcl11b* (encoding BCL11B), a transcriptional repressor key in maintenance of T cell lineage commitment and repression of NK differentiation (*37, 50*), was decreased in *Dot1L* KO CD8 T cells (Fig. 4D), likely contributing to the enriched NK gene signature (Fig. 4C). It should, however, be noted that most of the fold-changes are modest, indicating that it is likely a combined effect of multiple changes that drive the altered transcriptional state observed in the absence of DOT1L.

Combined, our findings show that DOT1L-deficient CD8 T cells lose certain aspects of their T cell identity, gain effector CD8 and NK cell features, and potentially acquire a rewired state in favor of increased cytolytic activity. Whether the mentioned transcriptional changes of specific genes in *Dot1L* KO T cells are directly mediated through the absence of DOT1L, rather than acquired through secondary effectors, still needs to be established.

To explore how the transcriptional changes due to loss of DOT1L correspond to changes at the protein level, whole-cell proteomics was performed on expanded CD8 T cells on Day 8. Although proteomics was not as sensitive as transcriptomics and was biased towards more abundant factors, several important observations were made from this analysis. In total, 282 proteins were upregulated and 113 downregulated in KO CD8 T cells (Fig. 4E). Overlaying the proteome changes and the transcriptome changes, indicated a similar expression trend (Fig. 4F-G). Importantly, among the most upregulated proteins in *Dot1L* KO, many cytotoxic granule proteins (granzymes and perforin, except Granzyme B) were found (Fig. 4E,H). This suggests that besides the increased degranulation upon exposure to target cells (Fig. 3I and Supplementary Fig. S3K), *Dot1L* KO cells have a higher intrinsic cytotoxic capacity. On the other hand, among the most downregulated proteins in KO were CD8 and CD5, confirming the surface staining results. However, TCR-associated CD3 ψ, ο, χ, and ζ (CD247) proteins were not uniformly altered in the whole-cell proteome of *Dot1L* KO T cells, nor at the transcriptional level (Fig. 4H-I). Their altered surface expression levels observed by flow cytometry (Fig. 2C and Supplementary Fig. S2B) might be explained in part by the reduced mRNA and protein expression of Cd247, encoding the CD3ζ chain, which is rate-limiting for the assembly and transfer of TCR/CD3 complexes on the cell surface (*51*). This would be in line with previous findings in the LcK-Cre;Dot1L model, in which *Dot1L* was deleted early in the T cell lineage (*4*). CD6 was also strongly downregulated at the protein level. This relatively understudied signal-transducing transmembrane receptor is expressed by T cells and some B and NK cells. Antibodies blocking the interaction between CD6 and its ligands augment anti-tumor immune responses though effects on CD8 T and NK cells (*52–54*). In agreement with the global correlations, the alterations in the expression levels of the selected proteins largely correlated with changes at the RNA level (Fig. 4H-I). Together, both transcriptome and proteome changes confirm a rewiring in *Dot1L* KO CD8 T cells towards a non-canonical intermediary NK and T cell phenotype.

### DOT1L affects CD8 T cell identity and function via its catalytic activity

DOT1L and its orthologue Dot1 in yeast have been shown to execute functions by methylation-dependent as well as methylation-independent mechanisms (*15, 16, 18–20*). To delineate whether the transcriptional role of DOT1L in CD8 T cells is linked to its methyltransferase activity, we took advantage of the availability of highly specific DOT1L inhibitors. Instead of adding 4-OHT, which genetically deletes *Dot1L*, the DOT1L inhibitor (DOT1Li) SGC-0946 was added during the *in vitro* activation and expansion of isolated OT-I CD8 T cells (Fig. 5A) (*55*). Treatment with SGC-0946, which inhibits DOT1L without affecting its protein expression level (*55*), slightly impaired the initial proliferative capacity of CD8 T cells during *in vitro* expansion (Fig. 5B). Like *Dot1L* KO, treatment with SGC-0946 did not affect H3K79me2 immediately but resulted in the loss of H3K79me2 over time and reached a level similar to *Dot1L* KO at Day 8 (Fig. 5C).

**Figure 5.**
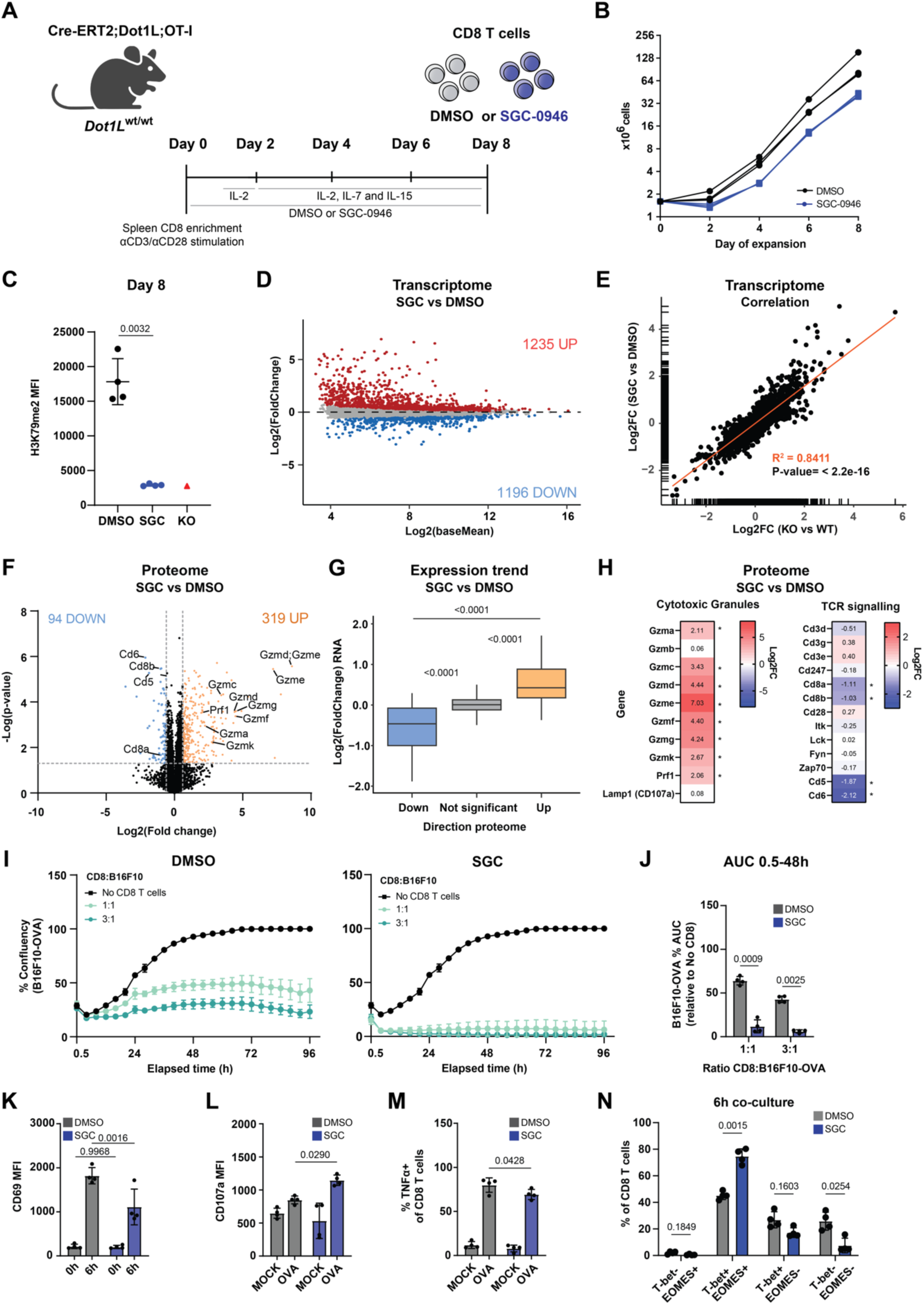
Loss of catalytic activity of DOT1L increases the killing potential of CD8 T cells. (A) Experimental set-up: mature OT-I CD8 T cells were treated with DOT1L inhibitor (10 µM SGC-0946) or DMSO (control) from Day 0 onwards. (B) Expansion of DOT1L inhibited and control CD8 T cells. (C) Flow cytometry-based quantification of H3K79me2 of DMSO and SGC-0946 treated CD8 T cells (n=4, paired treatment and expansion of CD8 T cells from individual mice). LCK-Cre *Dot1L* KO CD8 T cells (deletion in early T cell lineage *in vitro*) were used as a negative control (n=1). Significance was determined by a paired t-test. (D) MA plot showing differentially expressed genes between SGC-0946 and DMSO treated CD8 T cells. Red represents upregulated genes, and blue represents downregulated genes in in SGC-0946 treated CD8 T cells. (E) Correlation of differential gene expression for SGC-0946 vs DMSO and KO vs WT. The displayed correlation coefficient and significance was determined by Pearson’s product-moment correlation. (F) Volcano plot showing differentially expressed proteins between SGC-0946 and DMSO treated T cells. Orange represents upregulated proteins; blue represents downregulated proteins in SGC-0946 treated CD8 T cells. (G) Box plot showing the mRNA changes of proteins that were either differentially downregulated (blue), upregulated (yellow), or not significantly changed (grey) in SCG compared to DMSO treatment. (H) Heatmaps of selected proteins (see F) involved in cytotoxic granules and TCR signaling. Asterisk represents adjusted P-value (Padj) significance. (I) Quantification of the live cell imaging cytotoxic killing assay. Co-culture of DMSO or SGC-0946 treated CD8 T cells with B16F10-OVA target cells. The standard deviation represents variation between independent co-cultures with expanded CD8 T cells from different mice treated with either DMSO or SGC-0946 (paired). (J) Quantification of the area under the curve of I, relative to no CD8 (AUC at 48h), average +/− SD (n=4). (K) Flow cytometry surface expression of CD69 for CD8 T cells prior to co culture and after 6h co-culture with B16F10-OVA cells. (L) Flow cytometry expression of CD107a (degranulation) and (M) % TNFα producing CD8 T cells. MOCK represents B16F10 cells that do not express the OVA peptide, average +/− SD (n=4). (N) Flow cytometry quantification of T-bet and EOMES states after 6 hours of co-culture of SGC-0946 or DMSO treated CD8 T cells with B16F10-OVA cells (n=4, paired biological replicates). For J-N, significance was determined by paired two-way ANOVA and Šídák’s multiple comparisons tests.

Having established that H3K79me2 was lost, we subjected the DOT1L-inhibited and DMSO-treated control cells of Day 8 to transcriptomic and proteomic analyses. At the RNA level, 1235 genes were upregulated, and 1196 genes downregulated in SGC-0946 compared to DMSO treated CD8 T cells (Fig. 5D). Importantly, the differential gene expression pattern caused by DOT1L inhibition and *Dot1L* KO strongly correlated (Fig. 5E). Moreover, the differentially expressed genes in SGC-0946 treated cells showed the same gene set enrichments as *Dot1L* KO CD8 T cells, further strengthening this conclusion (Supplementary Fig. S5A-B). At the proteome level, 319 proteins were upregulated, and 94 proteins were downregulated upon SGC-0946 treatment (Fig. 5F) and the proteome changes correlated with their corresponding transcriptome changes (Fig. 5G; Supplementary Fig. S5C). Like the results from *Dot1L* KO, CD8, CD5 and CD6 proteins were downregulated, and multiple Granzymes and Perforin were upregulated upon SGC-0946 treatment (Fig. 5H, Supplementary Fig. S5E-G, and see the corresponding transcriptome changes in Supplementary Fig. S5C-D). These results reveal that the molecular changes in SGC-0946 treated and expanded CD8 T cells closely resemble those observed in *Dot1L* KO.

In line with these findings, SGC-0946 treated OT-I CD8 T cells showed enhanced target cell control *in vitro* compared to DMSO treated cells (Fig. 5I). Moreover, the killing was notably accelerated and resulted in near complete eradication of all target cells within 8 hours of co-culture (Fig. 5I-J, Supplementary Fig. S5H-I). SGC-0946 treatment by itself did not affect B16F10-OVA outgrowth (Supplementary Fig. S5J). During co-culture with target cells, SGC-0946 treated CD8 T cells showed reduced expression of the activation marker CD69 compared to DMSO treated cells, with an increased expression of the degranulation marker CD107a and decreased TNFα production, strongly resembling the phenotypes of *Dot1L* KO (Fig. 5K-M). Moreover, like *Dot1L* KO, SGC-0946 treatment did not compromise viability of CD8 T cells before or after 6h co-culture (Supplementary Fig. S5K). SGC-0946 treatment led to retention of a CD44^int^ CD62L^+^ and T-bet and EOMES double-positive state, suggesting a bias towards an intermediate state instead of a terminal differentiation state (Fig. 5N, Supplementary Fig. 5L-N). Combined, these findings suggest that DOT1L affects CD8 T cell identity and function at least largely via its catalytic activity.

### The catalytic function of DOT1L is dose-dependent and associated with H3K79 methylation

Following their discovery as epigenetic writers, many histone methyltransferases have been shown to also act on non-canonical substrates. Therefore, we investigated how the role of DOT1L in T cells relates to methylation of H3K79. Upon loss of DOT1L activity, H3K79me loss is slow and dependent on passive dilution mediated by cellular replication (Supplementary Fig. S6A, Fig. 1C) (*28, 29, 56, 57*). These slow kinetics severely hamper distinguishing early from late effects, and direct from indirect effects of H3K79me, respectively. However, gaining more mechanistic insight into these effects could be useful for potential targeted therapies in the future. Looking at when cellular changes occurred following loss of H3K79me, surface expression of CD3, CD8, CD5 and expression of EOMES were unaffected two days following treatment with DOT1L inhibitor SGC-0946 (H2K79me2 high) while all except EOMES were reduced after 8 days of expansion (H3K79me2 low; Supplementary Fig. S6A). These results suggest that manifestation of the molecular and phenotypic changes associated with the loss of DOT1L activity likely relate to the subsequent and progressive loss of H3K79 methylation due to passive dilution.

To further test this model in an independent way, we modified the experimental protocol to take advantage of the slow turnover of H3K79me. In this genetic approach, instead of deletion of *Dot1L in vitro*, Cre-ERT2;Dot1L^fl/fl^;OT-I mice were treated for three consecutive days with tamoxifen to induce deletion of *Dot1L in vivo* prior to T cell isolation. One day after the last treatment, denoted as Day 0, CD8 T cells were isolated and analyzed. While *Dot1L* deletion was efficient (Supplementary Fig. S6B-C), global H3K79me2 was not detectably altered in these cells (Fig. 6B and Supplementary Fig. S6D). Since DOT1L has been reported to have a relatively short half-life of approximately 2 hours (*58, 59*), this result supports the notion that upon loss of DOT1L protein, H3K79me2 loss in T cells is mainly dependent on replication-dependent dilution of post-translationally modified histones by incorporation of newly synthesized unmodified ones. Co-culture of these *Dot1L* KO H3K79me2+ CD8 T cells with B16F10-OVA-mCherry tumor cells revealed equally capable killing of target cells, compared to WT (Fig. 6C and Supplementary Fig. S6E-H). When these *Dot1L* KO H3K79me2+ cells were activated and expanded for 8 days *in vitro* and then tested, the *Dot1L* KO cells did lose H3K79me2 and showed enhanced tumor cell killing (Supplementary Fig. 6I-K), in line with what was observed following induction of *Dot1L* KO or DOT1L inhibition *in vitro*. Together, these findings support the idea that the loss of H3K79me is responsible for the molecular and cellular changes observed after *Dot1L* KO or DOT1Li (*e.g.* Fig. 3B-C & Fig. 5I-J). This indicates that the role of DOT1L in T cells is at least in part mediated via its canonical epigenetic writing function.

**Figure 6.**
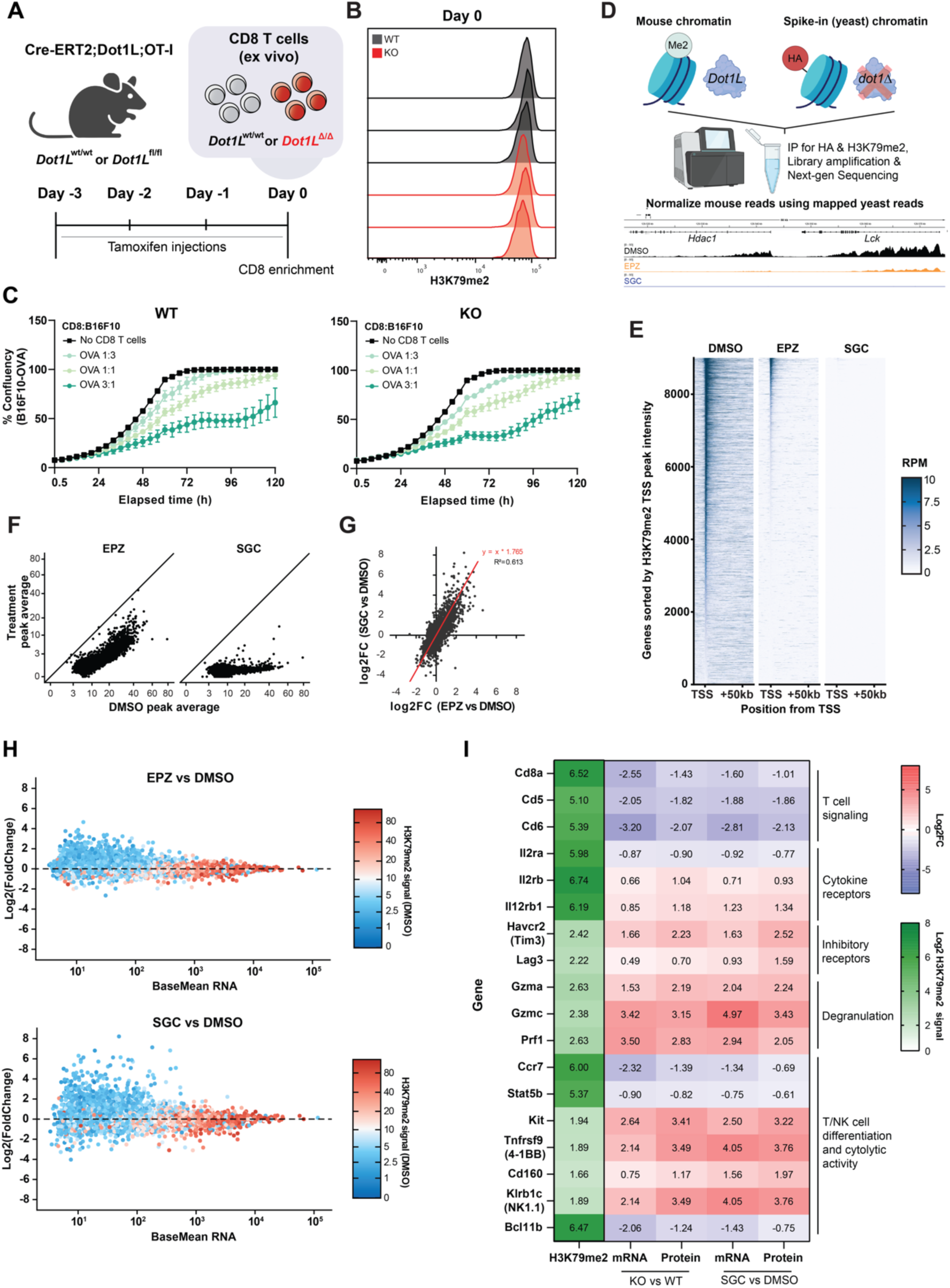
Loss of H3K79me2 is required to induce CD8 T cell phenotypes and NK-like differentiation. (A) Overview experimental set-up: RCM2;*Dot1L^wt/wt^*(WT) and *Dot1L^fl/fl^* (KO) mice (n=3) were treated with tamoxifen (75 mg/kg) for three consecutive days to genetically ablate *Dot1L*, after which mice were sacrificed, and splenic OT-I CD8 T cells were enriched. (B) Flow cytometry histogram of H3K79me2 of WT and KO mice at day 0 (n=3). H3K79me2 of enriched splenic CD8 T cells of individual mice displayed. (C) Quantification of live cell imaging of co-cultures with different ratios of CD8 to target cells (B16F10-OVA) (n=3) or no CD8. CD8s were added to the co-culture immediately following treatment and enrichment as described in (A). (D) Experimental set-up of spike-in normalized H3K79me2 ChIP-seq: sheared chromatin from HA-tagged *dot1Δ* yeast was co-immuno-precipitated with mouse chromatin samples to correct for global changes in H3K79me2. Representative tracks are shown for the *Lck* and *Hdac1* genes. (E) Tornado plot ranking genes based on H3K79me2 signal up to +2kb from the TSS. Data shown are aggregates of (n=2) IPs from CD8 T cells treated with DMSO (0.1%), EPZ-5676 (10 µM) or SGC-0946 (10 µM), after 8 days of expansion. (F) Scatterplot of H3K79me2 TSS peak average signal between DMSO (x-axis) and EPZ-5676 or SGC-0946 treated cells (y-axis). (G) Correlation of differential gene expression between EPZ-5676 and SGC-0946 treatment, relative to control (DMSO) (n=4). The displayed correlation coefficient and significance was determined by Pearson’s product-moment correlation. (H) MA-plots showing differentially expressed genes for EPZ-5676 or SGC-0946 relative to control (DMSO), overlaid with H3K79me2 signal in DMSO up to +2kb from the TSS (color gradient). (I) Heatmap displaying peak H3K79me2 TSS peak signal in DMSO (n=2; green gradient) and transcriptional and proteomic changes at selected genes in *Dot1L*-KO vs WT (n=4) and SGC vs DMSO treated (n=4) CD8 T cells (red-blue gradient). All data shown is significant based on adjusted P-values (Padj).

Even though distinguishing direct from indirect effects of loss of H3K79me2 is challenging, we performed ChIP-sequencing on T cells treated with DMSO (0.1 %, vehicle) or the DOT1L inhibitors SGC-0946 (10 µM) or EPZ-5676 (Pinometostat, 10 µM) for 8 days to gain more insight into how H3K79me and its loss might affect the identity of CD8 T cells. Treatment with SGC-0946 led to undetectable levels of H3K79me similar to those in KO, whereas EPZ-5676 resulted in partial loss of H3K79me (Supplementary Fig. S6L-M). While in previous studies higher doses of the structurally related DOT1L inhibitor EPZ004777 were used to inhibit DOT1L in T cells (*24*), we chose to use a maximum concentration of 10 µM as this has been commonly used as the highest dose and since higher concentrations have been reported to be toxic to cells that are otherwise known to be resistant to loss of DOT1L activity (e.g. see (*12, 25, 60*)).

To enable a comparative quantitative ChIP analysis of DOT1Li treated and untreated cells, we developed a spike-in protocol with an external reference that can be used for normalization independent of global changes in a sample. For that purpose, a yeast strain was generated that expresses HA-tagged H3 and in which Dot1 is deleted to eliminate all H3K79me. When mixed with mouse T cell chromatin, and when in addition to the anti-H3K79me antibody an anti-HA antibody is coupled to the beads, the yeast chromatin is enriched independently of the mouse H3K79me chromatin (Fig. 6D and Supplementary Fig. S6N-R). The spike-in approach is universal and convenient and given the ease of yeast engineering, it can be easily modified for application to other epigenetic marks. Using this external spike-in method, we performed ChIP for H3K79me2, because this methylation state shows a clear demarcation of 5’ ends of gene bodies and correlates well with transcriptional activity (see below) (*4, 6, 61–64*). When normalized to the yeast spike-in, SGC-0946 treated CD8 T cells showed near-complete loss of H3K79me2 ChIP signal across the genome, while EPZ-5676 showed a partial loss (Fig. 6D-F, Supplementary Fig. S6S-U). Inspection of the H3K79me2 levels around the transcription start sites (TSSs) of all protein-coding genes in EPZ-5676 treated cells showed a proportional loss of signal compared to that in DMSO treated cells and a reduction in the number of H3K79me2 marked TSSs largely without the gain of new ones (Fig. 6F and Supplementary Fig. S6S). This indicates that partial DOT1L inhibition affected all H3K79me2 demarcated genes in an equal manner and that DOT1L in expanding T cells does not have a subset of preferred target genes that maintain WT levels of H3K79me2 when DOT1L activity becomes limiting (Fig. 6F).

The global intermediate loss of H3K79me2 in EPZ-5676 treated cells compared to near complete loss upon SGC-0946 treatment correlated with intermediate effects on mRNA expression (Fig. 6G). Positive and negative changes in gene expression were observed in DOT1Li treated cells, and in agreement with the partial loss of H3K79me2 for EPZ-5676 treatment, the magnitude of changes was less pronounced in both compared to SGC-0946 treatment (correlation between both inhibitors R^2^=0.613, differential effect size 1.8-fold; Fig. 6G). Apparently, in CD8 T cells the loss of DOT1L affects the global H3K79me2 methylome and transcriptome in a dose-dependent manner. To investigate how DOT1L activity might affect gene expression, we first confirmed that H3K79me2 correlates with the level of gene transcription (*61, 65*) (Fig. 6H and Supplementary Fig. S6U, left panel). Partial DOT1L inhibition due to EPZ-5676 treatment led to partial and proportional loss of H3K79me2, with lowly expressed genes dropping to baseline levels, and highly expressed genes dropping to intermediate levels of H3K79me2 (Supplementary Fig. S6U middle panel). Interestingly, for the latter set of genes, lowering H3K79me2 to an intermediate level did not affect their expression indicating a threshold level of H3K79me2 required for maintaining gene expression.

Overlaying the H3K79me2 signals around the TSS in DMSO-treated cells onto the mRNA changes in DOT1Li vs DMSO treatment revealed that H3K79me2 was also more enriched on highly transcribed genes (high BaseMean) and nearly absent from lowly expressed genes (low BaseMean) (Fig. 6H). Since upregulation of gene expression was mostly observed for lowly expressed genes, which lacked H3K79me2, this suggests that in general the gene derepression effect observed upon DOT1L inhibition was most likely not causally related to loss of H3K79me2. Within the more highly expressed genes, the largest fold changes in expression were observed for genes that were downregulated following SGC-0946 treatment (Fig. 6H). This suggests that DOT1L’s main impact is likely through promoting gene expression of a subset of the genes that it methylates. At this point, it is unknown which of the genes that DOT1L methylates directly require DOT1L and H3K79me2 for their expression and which genes are regulated by indirect effects. However, our results do suggest that among all the differentially expressed genes, a relatively small subset is likely to be directly regulated by H3K79me2. Fig. 6I visualizes the relationship between the level of H3K79me2 in control CD8 T cells and the differential regulation at the RNA and protein level upon genetic ablation or chemical inhibition of DOT1L activity for a selected set of factors. Expression of the CD5/6/8 surface markers, the transcriptional repressor BCL11B and the cytokine receptor IL2RA are likely controlled by the catalytic activity of DOT1L and the high H3K79me2 levels of their corresponding genes suggests that they may be among the direct targets of DOT1L. On the other hand, the increased expression of Granzymes and Perforin is most likely caused by indirect effects because the genes harbor low levels of H3K79me2. A similar observation is found for certain molecules belonging to T/NK differentiation and cytolytic activity. While further studies are needed to deconstruct the mechanistic network underpinning the molecular and cellular changes, our data provide a valuable resource for untangling the downstream effects of DOT1L and for further tuning of T cells to harness the improved cell-intrinsic cytotoxic potential upon DOT1L targeting.

## Discussion

Deconstructing the role of epigenetic mechanisms in cytotoxic CD8 T cells is crucial to understand CD8 T cell cellular plasticity and differentiation trajectories and determine how epigenome perturbation can be exploited to modulate their immunological responses. The histone methyltransferase DOT1L is emerging as a key epigenetic and regulatory player in cellular differentiation, particularly in immune cells (*66, 67*). In various mouse models, the absence of DOT1L leads to altered cellular states, with a proposed role for DOT1L, and by inference H3K79me, in establishing an epigenetically based differentiation barrier to maintain cell identity. However, the cell-intrinsic role of DOT1L in CD8 T cell remains poorly defined (*4, 24, 25*).

Here we combined three different conditional models to establish the cell-intrinsic role of DOT1L in mature CD8 T cells in the mouse: deletion *in vitro*, inhibition *in vitro*, and deletion *in vivo*. Together, these findings suggest that DOT1L maintains the epigenetic and transcriptomic identity of CD8 T cells, prevents acquisition of NK-cell features, and restricts a cell-intrinsic cytotoxic tumor-cell killing potential. DOT1L does so via its catalytic activity and the loss-of-function phenotypes are correlated with loss of H3K79me, strongly supporting the idea that epigenetic mechanisms of gene regulation play a central role. DOT1L likely operates by nudging gene expression rather than by on-off switches because the changes in gene expression upon treatment with DOT1L inhibitor are relatively modest and dynamically dose-dependent. This also suggests that the altered cell state and function of CD8 T cells upon inactivation of DOT1L is likely mediated by a combinatorial effect of multiple changes. What remains to be further explored are the mechanisms by which blocking DOT1L rewires molecular and cellular networks and which of these components are the cause or consequence of this altered differentiation. The integrative analysis of transcriptome, proteome, and quantitative changes in H3K79me levels using a novel spike-in method provides an important framework for identifying indirect and candidate direct effects.

Upon early *in vivo* deletion of *Dot1L*, CD8 T cells differentiate prematurely towards a memory-phenotype state in an antigen-independent manner (*4*). A propensity to differentiate to memory-like cells would be compatible with the enhanced and accelerated cytotoxic response of *Dot1L* ablated mature T cells and the increased viability and survival of T cells after challenge with tumor cells. However, in our current *in vitro* deletion model, differentiation is mainly driven by the activation of T cells to allow for replication-mediated loss of H3K79me upon *Dot1L* deletion, resulting in effector-memory and memory like cell states. Nevertheless, in *Dot1L* KO, the transcriptome was positively enriched for genes annotated as ‘antigen response’ and analysis of differentiation markers indicated a trend of KO cells to attain an altered differentiation state, a finding similar to those made in the B cell lineage (*6, 7*).

Loss of *Dot1L in vitro* was well tolerated and upon adoptive cell transfer, *Dot1L* KO CD8 T cells were as proficient as WT CD8 T cells in controlling tumor outgrowth. Remarkably, *Dot1L* KO CD8 T cells showed enhanced tumor cell control *in vitro* despite the reduction in surface expression of TCR and CD8 prior to exposure to target cells and reduced expression of the activation marker CD69 upon challenge with target cells. Several factors might contribute to these phenotypic and functional changes. CD5, a negative regulator of TCR signaling (*32, 68*), is downregulated, which might compensate for the lower TCR and CD8 levels. Downregulation of CD6 might also be involved. CD6 has recently been proposed as a negative regulator in CD8 T cells and perturbing its interaction with ligands by blocking antibodies improves anti-tumor responses (*52, 54, 69*). Little is known about how CD6 is regulated but here we identify DOT1L as a factor that promotes CD6 expression. Another factor is that the high-avidity OT-I TCR (*70*) might compensate for the lower TCR expression to maintain the normal threshold for TCR stimulation. Of note, DOT1L inhibition has been shown to elevate the TCR stimulation threshold in human T cells, specifically in low-avidity T cell responses, by indirectly increasing the ERK phosphatase DUSP6 expression (*25*). The lower surface levels of TCR and CD8 that we observed previously as well as in this study likely contribute to reduced TCR stimulation upon DOT1L inhibition, especially for low-avidity TCRs.

Finally, regarding the differentiation state of *Dot1L* KO cells, the differential transcriptome in *Dot1L* KO cells was enriched for signatures associated with NK cells and T to NK cell differentiation. In line with this, *Dot1L* KO CD8 T cells showed increased expression of many Granzymes, Perforin, and the degranulation marker CD107a, which has been reported to be a marker of NK cell functional activity (*71*), although these latter factors are also induced in effector CD8 T cells. Granzyme B, differentially regulated between naïve and effector CD8 T cells and often used as marker for T cell functionality was not differentially expressed at the RNA or protein level in *Dot1L* KO cells. This suggests that loss of *Dot1L* in CD8 T cells initiates a non-canonical differentiation trajectory, which includes the upregulation of factors associated with NK cells. The acquisition of NK markers and NK-like cytolytic properties likely also contributes to the increased cytotoxic activity of CD8 T cells when DOT1L is ablated. Of note, activation of DOT1L-ablated OT-I CD8 T cells by target cells as well as their increased cytotoxicity towards tumor cells *in vitro* was still dependent on the TCR-antigen interaction (see MOCK controls in Supplementary Figs. S3I-L and S6K) indicating functional features of their T cell identity. Of note, DOT1L loss was recently shown to also control differentiation of *bona fide* NK cells (*10*).

T cells and NK cells cannot always be developmentally separated but may instead represent a continuum of cell fates. T cells and NK cells show many similarities in the way they kill target cells and how they differentiate upon activation (*72, 73*). Moreover, specific subsets of CD8 T cells express markers associated with NK cells and this has been associated with enhanced cytotoxic activity (*74, 75*). Recently, the dual blockade of PD-1 and LAG-3 in mouse CD8 T cells was shown to lead to expression of NK receptors and a stronger effector CD8 T cell program (*76–79*). NK cells are also emerging as promising targets for engineering, offering adoptive cell immunotherapy strategies that are complementary to and sometimes superior to those based on CD8 and CD4 T cells (*80, 81*). The various mechanisms underlying the induction of NK-like behavior in CD8 T cells and the implications of the observed molecular changes for the cellular activities are still incompletely understood (*73, 79*).

Our studies point to DOT1L as an epigenetic factor that warrants the identity and behavior of CD8 T cells and restricts the acquisition of NK-like features in CD8 T cells. Future studies will be aimed at determining the contribution of the various observed changes to the enhanced cytotoxic killing of *Dot1L* KO CD8 T cells. However, the net result is that loss of DOT1L activity in mature OT-I CD8 T cells improves their cytotoxic activity towards tumor cells in a cell-intrinsic manner *in vitro,* which is a strong starting point for exploring strategies for improvement of T cell responses by epigenetic perturbation.

## Material and Methods

### Mice

The derivation of Lck-Cre;*Dot1L*^fl/fl^;OT-I mice has been described previously (*4, 82, 83*). RCM2;*Dot1L*^fl/fl^ mice were derived by crossing Cre-ERT2 mice with *Dot1L*^fl/fl^ mice. Cre-ERT2 mice have been described elsewhere and were generated by inserting Cre fused to the human estrogen receptor (Cre-ERT2) into the ROSA-26 locus (*82*). *Dot1L*^fl/fl^ mice were derived from the Dot1Ltm1a(KOMP)Wtsi line generated by the Wellcome Trust Sanger Institute (WTSI) and obtained from the KOMP Repository (www.komp.org) (*83*). Cre-ERT2;*Dot1L*^fl/fl^;OT-I mice were generated by crossing Cre-ERT2;*Dot1L*^fl/fl^ mice with OT-I (B6J carrying the OT-I T cell receptor transgenes) mice (kindly gifted by the Ton Schumacher group (NKI), originally from Jackson labs). All strains were bred, and all experiments were performed in house, under specific pathogen free conditions. Mice used in experiments were between 6 weeks and 10 months old and gender and age matched between experimental groups. All experiments were approved by the Animal Ethics Committee of the NKI and performed in accordance with institutional, national and European guidelines for animal care and use.

### CD8 T cell enrichment and expansion

An overview of reagents used can be found in the supplementary information (Supplementary Table S1). Spleens were harvested and single cell suspensions were made by meshing cells through a 70 µm strainer (BD Falcon). Erylysis was performed by incubating the cells in red blood cell lysis buffer (0.15M NH_4_Cl, 10 mM KHCO_3_, 0.2 mM EDTA) for 3 min on ice. Cell suspensions were enriched for T cells by Pan T cell isolation (Miltenyi) or CD8 (Dynabeads untouched) isolation (Thermo). These enrichment procedures were performed according to manufacturer’s protocol. Prior to CD8 T cell expansion, 12-well plates were coated with aCD3 (5 μg/ml in PBS; BD) and incubated o/n at 4ᵒC. CD8 T cells were expanded by resuspension in whole T cell media (RPMI (Gibco), 10 % FBS heat-inactivated, 2 mM L-glutamine, 28.6 mM 2-mercaptoethanol, 1 mM sodium pyruvate, 10 mM HEPES, P/S) supplemented with aCD28 (22 μg/ml; BD) and subsequent plating on PBS-rinsed aCD3 pre-coated wells at 1-2 million cells per well in 2 ml and cultured at 37ᵒC with 5 % CO_2_. For Cre-ERT2 CD8 T cell experiments the media was further supplemented with 10nM 4-hydroxytamoxifen (4OHT; Sigma). One day after plating, IL-2 (Novartis) in whole T cell media was supplemented to a final concentration of 50 U/ml. From day 2 until day 8 (after plating), CD8 T cells were counted and resuspended in fresh whole T cell media supplemented with IL-2 (50 U/ml), IL-7 (5 ng/ml; Peprotech) and IL-15 (5 ng/ml; Peprotech), every other day. For inhibitor studies OT-I CD8 T cells were treated with 10 µM EPZ-5676 (Selleck), 10 µM SGC-0946 (Bio-Connect) or DMSO (0.1 %) from day 0. Resuspension in fresh cytokine supplemented T cell media with inhibitors or DMSO was performed every other day, when CD8 T cells were counted. For analysis of primary, *Dot1L*-KO non-expanded CD8 T cells, *Dot1L*-floxed and *Dot1L*-wild type Cre-ERT2 OT-I mice were treated with Tamoxifen (Sigma) in corn oil (20 mg/ml, dose 75mg/kg) for three consecutive days. 24h after the last treatment, mice were sacrificed, and splenic CD8 T cells were enriched using the Dynabeads™ Untouched™ Mouse CD8 Cells Kit according to manufacturer’s protocol.

### Generation of Lentivirus

An overview of all plasmids can be found in the supplementary information (Supplementary Table S1). The FlipGFP-T2A-mCherry gene insert (*84*) (Addgene #124428) was cloned into a lentiviral backbone (*85*) in order to create LentiV5-FlipGFP-Puro. H293T cells were transfected with LentiV5-FlipGFP-Puro, pLenti-dOVA(SIINFEKL)-Hygro or pLenti-MS2-P65-HSF1-Hygro control vector (OVA and MOCK control vector were kind gifts from the lab of Daniel Peeper, NKI), combined with lentiviral packaging plasmids pMD2G, pMDLg and pRSV-Rev (*86*) (Addgene #12259, #12251 and #12253) in DMEM supplemented with 10 % FBS and P/S and cultured at 37°C 5 % CO_2_. At 24 and 48 hours the supernatant was collected, 0.45µm filtered and stored at −80°C for later use in viral transductions.

### Generation and culture of mCherry positive B16F10-OVA and B16F10-MOCK control target cells

B16F10 cells (kind gift from the lab of Daniel Peeper, NKI) were cultured in DMEM (Gibco) supplemented with 10 % FBS and P/S at 37°C 5 % CO_2_. The cells were transduced o/n with lentivirus containing LentiV5-FlipGFP-Puro, followed by 2-day 1μg/ml Puromycin (Invivogen) selection. After Puromycin selection, clones were generated and measured for mCherry positivity by microscopy. Selected clones were subsequently transduced o/n with lentivirus containing Lenti-dOVA-Hygro or the control insert, followed by seven days 200 μg/ml Hygromycin B (Life-technologies) selection. FlipGFP-OVA pools were treated for 48h with 5 μg/ml IFNy, gently treated with accutase for 2 minutes, collected, washed with PBS, stained for anti-SIINFEKL-H2Kb (Biolegend 141605) and FACS sorted on the FACSAria Fusion (BD Biosciences) for high OVA peptide presentation on H2K^b^.

### Live cell imaging cytotoxicity assay

Prior to seeding, B16F10 cells were treated for 48h with 5 μg/ml IFNy to upregulate MHC-I expression (*87*). Subsequently, cells were gently detached by treatment with Accutase for 2 minutes at 37°C and collected by rinsing with DMEM (Gibco) supplemented with 10 % FBS and P/S. A total of 7,500 B16F10-OVA cells were seeded per well in 96 wells µClear plates (Greiner) and incubated at 37°C 5 % CO_2_ for 4-5 hours in target cell media. Thirty minutes prior to imaging, supernatant was aspirated and harvested CD8 T cells in T cell media supplemented with IL-2 (50 U/ml; Novartis) were added in the corresponding numbers and ratios to the target cells. Live cell imaging was thereafter performed using the Incucyte ZOOM imaging system. Brightfield and mCherry images were obtained every 4 hours using the following settings: red 800ms, 1392 × 1040 at 1.22µm/pixel, Nikon 10x objective, dual color model 4459 filter module. After acquisition, time-course movies were generated and Brightfield and mCherry channel specific images were exported using the Incucyte ZOOM software (TIFF file format). The viable cell confluence over time was quantified using FiJi (ImageJ 2.3.0/1.53q / Java 1.8.0_322) and a custom build script optimized for quantification of mCherry from exported images. All images of a plate were used to generate a background subtraction file, Sigma of Gaussian blur was set on 3, Threshold bias at 0.5, opening iterations at 2 and dilation iterations at 2. Quantified data was plotted and the area under the curve (AUC) was quantified for statistical comparisons for killing capacity.

### Fixed time-point cytotoxicity assay

B16F10 cells were treated with IFNy and collected as described above. 70,000 B16F10-OVA cells were plated in 12 wells plates (Greiner) and incubated at 37°C, 5 % CO_2_ for 4-5 hours in DMEM (Gibco) supplemented with 10 % FBS and P/S. Media was aspirated and expanded CD8 T were added to the target cells in corresponding ratios to target cells in complete T cell media supplemented with 50 U/ml IL-2. After either 6h or 48h co-culture at 37°C, 5 % CO_2_, CD8 T cells were carefully collected and used in flow cytometry experiments. For 48h incubated co-cultures, target cells were fixed on the plate using 4 % formaldehyde (diluted from 37 %; Honeywell) in PBS for at least 30 minutes and subsequently stained in crystal violet staining solution (0.5 %; in MilliQ 20 % methanol) for 20 minutes at room temperature. After staining, the wells were washed three times by gently submerging the plate in water and air dried for 2-3 days. The plates were subsequently imaged using the Epson Perfection V750 PRO scanner (dual lid, 24-bit color, 300dpi).

Degranulation and cytokine production by CD8 T cells was determined by co-culturing B16F10-OVA cells with expanded CD8 T cells as described above for 5h, in presence of GolgiStop (BD), Monensin (Biolegend) and CD107a (1:200) as previously described (*34*). CD8 T cells were collected by careful pipetting and subsequently stained for flow cytometry as described above.

### *In vivo* tumor control

Recipient CD45.1 mice, female and aged 8-12 weeks, were injected subcutaneously with 200,000 B16F10-OVA tumor cells in PBS/Matrigel (1:1; 100µl) in one flank. At Day 7 after injection of tumor cells all mice were subjected to 5Gy total body irradiation (SmART+ irradiator (Precision)). At Day 8 recipient mice received either HBSS vehicle or 4 million *in vitro* expanded *Dot1L*-kockout (KO) or wild-type (WT) OT-I CD8 T cells (pooled cells from 4 female donor mice, 8 recipient mice per arm, age matched groups, blinded and cage randomized). After adoptive cell transfer, the mice were treated three consecutive days (Day 8-10) with 100,000 IU IL-2 (Novartis) by intraperitoneal injections. Tumor size was monitored, and tail vein punctures were taken over time to determine the presence and quantity of adoptively transferred OT-I CD8 T cells. At the endpoint, which was either welfare related or after reaching a tumor size of 1000 mm3, mice were sacrificed and tumor, spleen and blood samples were used for flow cytometric analysis. Spleen and blood samples were erylyzed on ice (3 minutes and 30 minutes, respectively). Mice that died prematurely were excluded from sample work-up but included in tumor growth and survival plots if they developed tumors of near endpoint size. Mice that suffered welfare related issues with near endpoint size tumors were not excluded from the analysis. Mice that did not develop a tumor were excluded from the analysis in its entirety.

Leukocytes were enriched by mashing the tumors through a 70 µm filter. Filters were rinsed with media and 10ml Ficoll (Sigma) was carefully added in the bottom of the 50ml tube containing the mashed cell suspension, bringing the total around 35-40ml. The tubes were centrifuged for 20 minutes at 536 x*g* at 4°C without brakes. The leukocyte-rich interphase was collected and used for staining. 2 million splenocytes and all remaining blood and tumor enriched leukocytes were stained for flow cytometric analysis. Tumor enriched leukocyte samples derived with less than 150 cells in their respective parent gate were excluded from analysis.

### Flow cytometry

Enriched, expanded or co-cultured CD8 T cell single cell suspensions (typically 200,000 cells per stain) were washed with PBS and subsequently stained with Zombie-NIR in PBS for 30 minutes on ice. Cells were pelleted and subsequently fluorescently labeled by antibodies in FACS buffer (0.5 % BSA and 2 mM EDTA in PBS) at dilutions indicated (Supplementary Table S1). Intracellular staining was performed according to manufacturer’s protocol, using the Transcription Factor Buffer kit (BD Biosciences). H3K79me2 staining was performed according to the method described previously (*4*). Briefly, after cell permeabilization, cells were resuspended in Transcription Factor Buffer kit Perm/Wash buffer supplemented with 0.25 % Sodium Dodecyl Sulfate (SDS) and stained with anti-H3K79me2 (Millipore) followed by 1:1000 secondary antibody staining (Goat-anti-Rabbit AF488; Invitrogen) in Perm/Wash. Annexin V (AnnV) and DAPI staining was performed by staining 50,000-100,000 CD8 T cells in 50 µL AnnV binding buffer (10 mM HEPES (pH7.4), 0.14M NaCl and 2.5 mM CaCl2, freshly prepared from 10x stock) at indicated dilutions of AnnV and DAPI (Supplementary Table S1) for 20 minutes at room temperature (dark). Subsequently the samples were diluted 3x in binding buffer and directly measured. Flow cytometry was performed using the LSR-Fortessa II (BD Biosciences). Data was analyzed using FlowJo V10.9.0 (BD Biosciences). Enriched CD8 T cells with aberrant phenotypes at Day 0, e.g. phenotypes suggesting unintended immune challenge or genetic aberrations in our Specific Pathogen Free (SPF) housed mice, were excluded from further analysis.

### RNAseq sample preparation

CD8 T cells were activated and expanded for 8 days as described above in whole T cell media. 1×10^6^ cells were transferred to RNase/DNase-free Eppendorf tubes, pelleted at 500 x*g* for 5 minutes, washed in 750 µl PBS, centrifuged for 10 minutes at 600 x g at 4ᵒC and resuspended in RLT buffer (Qiagen) supplemented with 1 % 2-mercaptoethanol. Samples were stored at −80ᵒC until further use.

### RNAseq sample work-up and library preparation

Total RNA was isolated according to manufacturer’s protocol using the RNAeasy mini kit (Qiagen). Quality and quantity control was performed on the 2100 Bioanalyzer on the Agilent RNA nanochip. For *Dot1L* KO and WT and EPZ-5676 (10 µM) and DMSO (0.1 %) treated CD8 T cells, RNA samples with an RNA Integrity Number >8 were subjected to library preparation. For SGC-0946 (10 µM) and DMSO (0.1 %) treated CD8 T cells, the RNA samples with an integrity number >6 were subjected to library preparation. Strand specific cDNA libraries were generated using the TruSeq Stranded mRNA sample preparation kit (Illumina) according to the manufacturer’s protocol. The libraries were analyzed for size and quantity of cDNAs on a 2100 Bioanalyzer using a DNA 7500 chip (Agilent), diluted, and pooled in multiplex sequencing pools. The paired-end 51 bp reads were sequenced using the NovaSeq 6000.

### RNAseq preprocessing

Strand specific RNA reads (11–33 million reads per sample), 51 bp paired-end, were aligned against the mouse reference genome (Ensembl build 38) using Tophat (version 2.1, bowtie version 1.1). Tophat was supplied with a Gene Transfer File (GTF, Ensembl version 77) and was supplied with the following parameters: “prefilter-multihits –no-coverage-search – bowtie1 –library-type fr-firststrand”. To count the number of reads per gene, a custom script which is based on the same ideas as HTSeq-count has been used. A list of the total number of uniquely mapped reads for each gene that is present in the provided Gene Transfer Format (GTF) file was generated.

### RNAseq data analysis

Subsequent analysis was performed in R version 4.2.2 with Bioconductor packages from release 3.16. Genes that have no expression across all samples within the dataset were filtered out. Data was pre-filtered and the analysis was restricted to the genes that have at least 20 counts in at least 4 samples to exclude very low abundance genes. Principal component analysis (PCA) was performed using the ‘prcomp’ function on variance stabilizing transformed data with the ‘vst’ function from the DESeq2 package (1.38.3) using default arguments and plotted by using ggplot2 package (3.5.0). Sample to sample distances obtained using ‘dist’ function on variance stabilizing transformed data were subjected to hierarchical cluster analysis and dendrogram preparation. Hierarchical cluster analysis for the samples was performed using ‘hclust’ function with default arguments. The dendrogram was made by using ‘dendro_data’ function from ggdendro package (0.2.0). Normalized counts from DESeqDataSet from the DESeq2 package were subjected to calculate correlation among the samples by using ‘cor’ function using spearman method. Differential gene expression analysis was performed using DESeq2 package and default arguments with the design set either according to the genotypes (WT and *Dot1L* KO) or treatment conditions (DMSO and SGC-0946) and plotted by using ggplot2 package (3.5.0). Adaptive effect size shrinkage was performed with the ashr package version 2.2.63 to control for the lower information content in low abundance transcripts. Genes were considered to be differentially expressed when the p-value of the negative binomial Wald test was below 0.01 after the Benjamini-Hochberg multiple testing correction. Pearson correlation between log2FoldChanges from different comparisons (KO vs WT and SGC vs DMSO) was calculated by using ‘cor.test’ function and plotted by using ‘ggscatter’ function under ggpubr package (0.6.0).

For the effect size comparison between EPZ-5676 and SGC-0946 treatments, normalized count tables for EPZ and SGC treatment were merged retaining the intersection of genes. For the symmetric difference set of genes, we confirmed low expression. To filter out genes with low expression, genes that did not have more than 10 counts in at least 4 samples were removed for further analysis. The DESeq2 R/Bioconductor package was used to test for differential expression using ‘∼ batch + treatment’ as design argument and otherwise default argument. Results from a negative binomial Wald test were retrieved with ‘contrast = c(“treatment”, “EPZ”/”SGC”, “DMSO”), alpha = 0.05, lfcThreshold = 0.1’. A gene was considered to be differentially expressed when the FDR corrected p-value was below 0.05. To compare effect sizes between EPZ and SGC treatment, a total least squares coefficient was obtained using the ‘prcomp’ R function.

### Gene set enrichment analysis

Gene set enrichment analysis (GSEA) was performed using the software provided by the UC San Diego and Broad institute (*88*) in combination with v2022.1.Mm of the Molecular Signatures Database (msigdb) and v2022.1.Mm for symbols. Gene Ontology (GO) analysis performed in GSEA includes molecular function, biological processes and cellular components. Phenotype permutation was set at 1000 times and the weighted signal to noise statistical approach was implemented to rank genes. For this study we focused on biological processes and selected features based on a nominal p-value <0.05 and FDR q-value <0.25 and relevance to CD8 T cell function. Moreover GSEA-based curated gene set enrichment was performed to compare enrichment in DOT1L ablated or inhibited cells with documented cell types and cellular states. Similarly, a cut-off was set at nominal p-value <0.05 and FDR q-value <0.25.

### Proteomics

CD8 T cells were activated and expanded for 8 days in whole T cell media as described above. For protein digestion, snap-frozen cell pellets (2.5 × 10^6^ cells; kept at −80ᵒC until used) were lysed in 5 % SDS lysis buffer, boiled and sonicated. Aliquots corresponding to 50 µg of protein were digested using S-Trap micro-columns (ProtiFi) according to the manufacturer’s protocol. In short, samples were reduced and alkylated using DTT (20 mM, 15 min, 55 C) and IAA (40 mM, 10 min). The samples were acidified and a methanol TEAB buffer was added, prior to loading on the S-Trap column. Trapped proteins were washed 4 times with the methanol TEAB buffer and then digested for 2 h at 47 C using Trypsin (Sigma). Digested peptides were eluted and dried in a vacuum centrifuge before LC-MS analysis.

For *Dot1L* WT and KO CD8 T cells: prior to mass spectrometry analysis, the peptides were reconstituted in 2 % formic acid. Peptide mixtures were analyzed by nano LC-MS/MS on an Orbitrap Exploris 480 Mass Spectrometer equipped with an EASY-NLC 1200 system (Thermo Scientific). Samples were directly loaded onto the analytical column (ReproSil-Pur 120 C18-AQ, 2.4 μm, 75 μm × 500 mm, packed in-house). Solvent A was 0.1 % formic acid/water and solvent B was 0.1 % formic acid/80 % acetonitrile. Samples were eluted from the analytical column at a constant flow of 250 nl/min in a 90-min gradient, containing a 78-min linear increase from 6 % to 30 % solvent B, followed by a 12-min wash at 90 % solvent B. The Exploris 480 was run in data-independent acquisition (DIA) mode, as described above.

For DMSO and SGC-0946 treated CD8 T cells: Prior to mass spectrometry analysis, the peptides were reconstituted in 0.1 % formic acid and loaded on the Evotip Pure™ (Evosep). Peptide mixtures were analyzed by nano LC-MS/MS on an Orbitrap Exploris 480 Mass Spectrometer equipped with an Evosep One LC system. Peptides were separated using the pre-programmed gradient (Extended method, 88 min gradient) on an EV1137 (Evosep) column with an EV1086 (Evosep) emitter. The Exploris 480 was run in data-independent acquisition (DIA) mode, with full MS resolution set to 120,000 at m/z 200, MS1 mass range was set from 350-1400, normalized AGC target was 300 % and maximum IT was 45ms. DIA was performed on precursors from 400-1000 in 48 windows of 13.5 Da with an overlap of 1 Da. Resolution was set to 30,000 and normalized CE was 27.

Raw data were analyzed by DIA-NN (version 1.8) (*89*) without a spectral library and with “Deep learning” option enabled. The Swissprot Mus Musculus database (17,082 entries, release 2021_07) was added for the library-free search. The Quantification strategy was set to Robust LC (high accuracy) and MBR option was enabled. The other settings were kept at the default values. Protein group abundances were extracted from the DIA-NN result files, imported into Perseus (version 2.0.7.0) (*90*) and Log2-transformed. Values were filtered for presence in at least 3 out of 4 replicates in at least one condition. Missing values were replaced by imputation based on a normal distribution (width: 0.3 and downshift: 2.4). Differentially expressed proteins were determined using a Student’s t-test (minimal threshold: FDR = 0.05 and S0 = 0.1).

Box plots showing the mRNA changes of three groups of proteins that were differentially downregulated, upregulated, or not significantly changed, were generated using ggplot2 package (3.5.0). The inter quartile range (IQR) hinges correspond to first and third quartiles, whiskers show 1.5 IQR from upper and lower hinge and the central horizontal line within the box represents the median value of the Log2 of fold change in RNA expression. Significance was determined using Wilcoxon pair wise statistical analysis.

### H3K79 methylation mass spectrometry sample preparation and data analysis

To measure H3K79 methylation levels, samples were prepared and analyzed in the same way as described previously (*91*) with the following exceptions. Peptide mixtures were analyzed by nano LC-MS/MS on a Thermo Orbitrap Fusion hybrid mass spectrometer (Thermo Scientific) equipped with an EASY-NLC 1200 system (Thermo Scientific). Samples were directly loaded onto the analytical column (ReproSil-Pur 120 C18-AQ, 2.4 μm, 75 μm × 500 mm, packed in-house). Solvent A was 0.1 % formic acid/water and solvent B was 0.1 % formic acid/80 % acetonitrile. Samples were eluted from the analytical column at a constant flow of 250 nl/min in a 45-min gradient, containing a 30-min linear increase from 10 % to 50 % solvent B, followed by a 15-min wash at 90 % solvent B. Nanospray was achieved using the Nanospray FlexTM Ion source with a liquid junction set-up at 1.95 kV. The instrument was run in top speed mode with 2s cycles. Parameters for quantification of the different methylation states were optimized using a set of four purified synthetic peptides with the sequence EIAQDFK*TDLR (K*: K-me0, -me1, -me2 and -me3), as described previously (*91*).

### ChIP-seq sample preparation

After 8 days of expansion in DMSO, EPZ-5676 or SGC-0946 containing media (final concentration 10 µM), 2.5× 10^6^ CD8 T cells were cross-linked in whole T cell media supplemented with 1 % methanol free formaldehyde (Thermo Scientific) by incubating at room temperature for 10 minutes under gentle agitation. Cross-linking was neutralized by adding glycine (final concentration 125 mM) and incubating for 5 minutes. The fixed samples were centrifuged for 5 minutes at 600 x*g*, washed once using PBS + protease inhibitor cocktail (PIC; Roche), and re-suspended in PBS + PIC + 10 % FBS. Samples then were transferred to RNase/DNase-free Eppendorf tubes, centrifuged for 5 minutes at 600 x*g*, aspirated, snap-frozen and stored at −80°C till subsequent use.

### ChIP-seq yeast spike-in preparation

To normalize all ChIP-reads, a yeast spike-in strain was generated, in which one gene copy encoding for histone H3 (HHT2) was genomically tagged with 3xHA (using plasmid pFvL160; Addgene #64753), and *DOT1* was replaced with a NatMX selection cassette. This yeast strain (NKI2575; MATa lys2Δ0 trp1Δ63 his3Δ200 ade2Δ::hisG ura3Δ0 leu2Δ0 met15Δ0 ADE2-TEL-VR URA3-TEL-VIIL dot1Δ::NATMX set1Δ::KANMX hht2::HHT2-LoxP-3xHA-HYG-LoxP-3xT7) was cultured in standard yeast media (YEPD) at 30°C to an OD660 of 0.5 prior to cross-linking. Cross-linking was performed by adding 10 % v/v fix solution (50 mM HEPES-KOH, pH 7.5, 100 mM NaCl, 1 mM EDTA, 11 % formaldehyde (Thermo Scientific)) and incubating at room temperature under gentle agitation for 20 minutes, after which glycine (final concentration 125 mM) was added for 5 minutes to quench the cross-linking. The fixed cells were centrifuged for 5 minutes at 4°C, 4000 x*g*, washed once using ice-cold TBS, and re-suspended in TBS+PMSF (Roche) +10 % FBS. The samples were then transferred to Eppendorf tubes, snap-frozen and stored at −80°C as described above.

From these samples, chromatin was isolated by re-suspending the pellet in ice-cold breaking buffer (100 mM TRIS pH 7.9, 20 % glycerol + PIC) with silica/zirconia beads (0.5 mm; BioSpec products) and bead-beating for 2x 2 minutes using the Mini-Beadbeater (Biospec products). Samples were then homogenized with FA buffer (50 mM HEPES-KOH, pH 7.5, 140 mM NaCl, 1 mM EDTA, 1 % Triton-X-100, 0.1 % Na-deoxycholate + PIC) and transferred to new RNase/DNase-free Eppendorf tubes. Chromatin was pelleted by centrifugation at 13000 xg for 1 minute, washed once using FA buffer + 0.13 % SDS, and resuspended in FA buffer + 0.8 % SDS. Samples were sonicated using the BioRuptor (PICO) for 6 cycles of 30 seconds on, 30 seconds off, centrifuged for 5 minutes at 13000 xg, re-suspended in FA buffer and stored at −80°C. An aliquot was subsequently thawed, treated with 10 mg/mL RNase A (Sigma) and 10 mg/mL Proteinase K (Sigma) and incubated for 1 hour at 50°C, then overnight at 65°C to reverse cross-links. DNA was eluted using a PCR purification kit (Qiagen) according to the manufacturer’s protocol. Eluted DNA was checked for shearing efficacy by running the samples through a 1 % agarose gel, and DNA concentration was determined by using the Qubit™ dsDNA Quantification Assay Kit (Thermo Scientific).

### Normalized H3K79me2 ChIP-seq sample work-up and library preparation

Cross-linked CD8 T cells were washed once with PBS + PIC, centrifuged for 5 minutes at 600 xg, re-suspended in ice-cold nuclei lysis buffer (50 mM Tris-HCl pH8, 10 mM EDTA pH8, 1 % SDS + PIC) and incubated on ice for 10 minutes. Samples were then sonicated for 3 minutes (3x 30 seconds on followed by 30 seconds off), after which debris was pelleted by centrifugation for 10 minutes at 4°C, 13000 xg. The supernatant was transferred to a new RNase/DNase-free Eppendorf tube and supplemented with sample buffer (9 parts ChIP dilution buffer (50 mM Tris-HCl pH 8, 0.167 M NaCl, 1.1 % Triton X-100, 0.11 % sodium deoxycholate) and 5 parts RIPA-150 (50 mM Tris-HCl pH8, 0.15M NaCl, 1 mM EDTA pH8, 0.1 % SDS, 1 % Triton X-100, 0.1 % sodium deoxycholate + PIC) per 1 part sample. Shearing efficiency was confirmed by transferring 10 % of the suspension to a new tube, and reversing the cross-links, eluting the DNA and running a gel and Qubit analysis as described above.

Based on Qubit DNA quantifications, the yeast chromatin spike-in was added 1:1000. Samples were homogenized and diluted in sample buffer, after which 10 % was removed and stored at −20°C as input. To the remaining sample, 0.5 μg mouse anti-HA antibody (manufactured at NKI) and 1 µl rabbit anti-H3K79me2 (Millipore) was added, after which samples were incubated under rotation at 4C overnight. Following incubation, Dynabeads™ Protein G (Thermo Scientific) in sample buffer were added, and samples were incubated under rotation at 4°C for 2 hours. The beads were then washed using the DynaMag-2 magnetic rack (Thermo Scientific), by aspirating the supernatant and adding 800 µl RIPA-150, followed by incubation under rotation at 4°C for 5 minutes. The washing steps were repeated twice using RIPA-500 (50 mM Tris-HCl pH 8, 0.5 M NaCl, 1 mM EDTA pH8, 0.1 % SDS, 1 % Triton X-100, 0.1 % sodium deoxycholate), twice using RIPA-LiCl (50 mM Tris-HCl pH 8, 1 mM EDTA pH 8, 1 % Nonidet P-40, 0.7 % sodium deoxycholate, 0.5 M LiCl2) and once using TE. Subsequently, samples were re-suspended in 150 µl Direct Elution buffer (10 mM Tris-HCl pH 8, 0.3 M NaCl, 5 mM EDTA pH8, 0.5 % SDS). Samples were de-cross-linked, quantified and quality checked as described above. All samples were library amplified and sequenced using the NovaSeq with paired end 51 bp reads.

### ChIP-seq data processing

ChIP-seq samples were mapped to a hybrid genome made from concatenating the mouse mm10 genome (Ensembl GRCm38) with the yeast sacCer3 genome. Mapping was performed using BWA-MEM (version 0.7.17-r1188) with the option ‘-M’ and filtered for a mapping quality of 10. Duplicate alignments were removed using MarkDuplicates from the Picard toolset (version 3.0.0) with the option ‘-REMOVE_DUPLICATES true’. Paired-end reads were then converted to fragments by reading in mapped, properly paired reads using the GenomicAlignments R/Bioconductor package (release 3.17). Peaks were called using MACS2 version 2.2.7.1 using samples pooled by treatment and pooled input samples as the ‘control’ argument using the options ‘-f BEDPE -g mm --keep-dup all --broad’. Peaks detected on yeast chromosomes were removed. Genomic bigwig tracks were generated for both individual samples and samples pooled by treatment from fragments mapping to the mouse genome using a custom R script using the ‘coverage’ function from the GenomicRanges R/Bioconductor package. The track heights were multiplied by a factor ‘1⋅10^6^ / number of yeast reads’ for values to represent ‘coverage per million yeast reads’ to allow quantitative comparisons. Signal values per peak were obtained by using the BigWigFile-method of the ‘summary’ function in the rtracklayer R/Bioconductor package. For aggregate visualization of coverage data, the tornadoplot R/GitHub package was used and TSS/exons were determined using the EnsDb.Mmusculus.v75 R/Bioconductor package. To calculate distances to the first internal exon, exons within a gene were first collapsed to disjoint ranges using the ‘reduce’ function from the GenomicRanges R/Bioconductor package. The distance was then taken as the number of basepairs between the 3’ end of the most 5’ exon and the 5’ end of the subsequent exon for genes that have at least 2 distinct exons.

### PCR

Enriched CD8 T cells were lysed using DirectPCR Lysis Reagent (mouse tail) (Viagen Biotech) with 1mg/ml proteinase K (Sigma). *Dot1L* WT, floxed, and deleted (exon 2) alleles were detected by PCR using the following primers, as described in Kwesi-Maliepaard, Aslam, Alemdehy, van den Brand, McLean, Vlaming, van Welsem, Korthout, Lancini, Hendriks, Ahrends, van Dinther, den Haan, Borst, de Wit, van Leeuwen and Jacobs (*4*). Dot1L_FWD: GCAAGCCTACAGCCTTCATC, Dot1L_REV: CACCGGATAGTCTCAATAATCTCA and Dot1L_Δ: GAACCACAGGATGCTTCAG. PCR reactions were performed using MyTaq Red Mix (GC Biotech). Agarose gel electrophoresis was performed to determine the genotype. Where relevant, the efficacy of floxing out exon 2 was determined by quantifying band densities using ImageJ (version 1.54g), correcting for product size and calculating the percentage by dividing the band density of the floxed-out band over the total band density within the sample.

### Visualization and statistical analysis

Data was visualized and statistical analyses were performed in Graphpad Prism 9, unless otherwise indicated. Paired or unpaired statistical analysis was applied according to the experimental set-up used. Subtests included all groups of a specific comparison, relevant p-values were plotted. The graphical abstract and subfigures 1A, 2A, 5A, 6A and 6D use images derived from BioRender, and were generated by using BioRender.com.

## Acknowledgements

We thank Julia Busselaar and Elselien Frijlink for discussion and valuable advice, David W. Vreedevoogd and Daniel Peeper for providing B16F10 WT cells and OVA and control plasmids, Martijn van Baalen, Debajit Bhowmick and Frank van Diepen for Flow cytometry advice and technical support, Thom Molenaar and Hans Teunissen for advice on ChIP, Ben Morris, Antonio Mulero Sánchez and Johan Kuiken for facilitating use of the IncuCyte® zoom and for gifted material for the experiments, Onno Bleijerveld for valuable advice on proteomics sampling, Wim Brugman and Roel Kluin for support with RNA sequencing and mapping to reference genomes and Robin van der Weide for mapping initial pilot experiment spike-in normalized ChIP-seq data, and the Research High Performance Computing (RHPC) facility of the Netherlands Cancer Institute (NKI) for hosting all data analyses We would further like to thank our colleagues from the Preclinical Intervention Unit of the Mouse Clinic for Cancer and Ageing (MCCA) at the NKI for their technical support performing the animal experiments. We thank Marianne Bakker and Marlize van Breugel for critically reading our manuscript. We thank the NIH Tetramer Core Facility (contract number 75N93020D00005) for providing (H-2K(b) SIINFEKL PE and H-2K(b) SIINFEKL BV421) tetramers.

## Funding & Statements

This research was supported by an institutional grant of the Dutch Cancer Society (KWF) and of the Dutch Ministry of Health, Welfare and Sports; by grants of the Dutch Cancer Society (KWF; NKI2018-1/11490 and 2022-2 EXPL/14479 to FvL and HJ) and by a grant from the ZonMW (TOP91218022 to FvL and HJ). LH and the proteomics facility are supported by the Dutch NWO X-omics Initiative.

The authors declare no competing interests.

## Author contributions

Conceptualization: MM, EMK-M, HJ, FvL

Validation: MM, WdL, MAA, EMK-M

Methodology, formal analysis and investigation: MM, WdL, EMK-M, MAA, MK, TvdB, LH, TvW, NP

Software: MAA, MM, BvdB, TvdB

Writing original draft: MM, WdL, HJ, FvL

Supervision: EdW, JB; HJ, FvL

Funding acquisition: HJ, FvL

## Notes

### Competing Interest Statement

The authors have declared no competing interest.

