## Supplementary Figures for "Histone methyltransferase DOT1L maintains identity and restricts cytotoxic potential of CD8 T cells"

**Document S1**. Supplementary Figures S1–S6

### Supplementary Figure S1

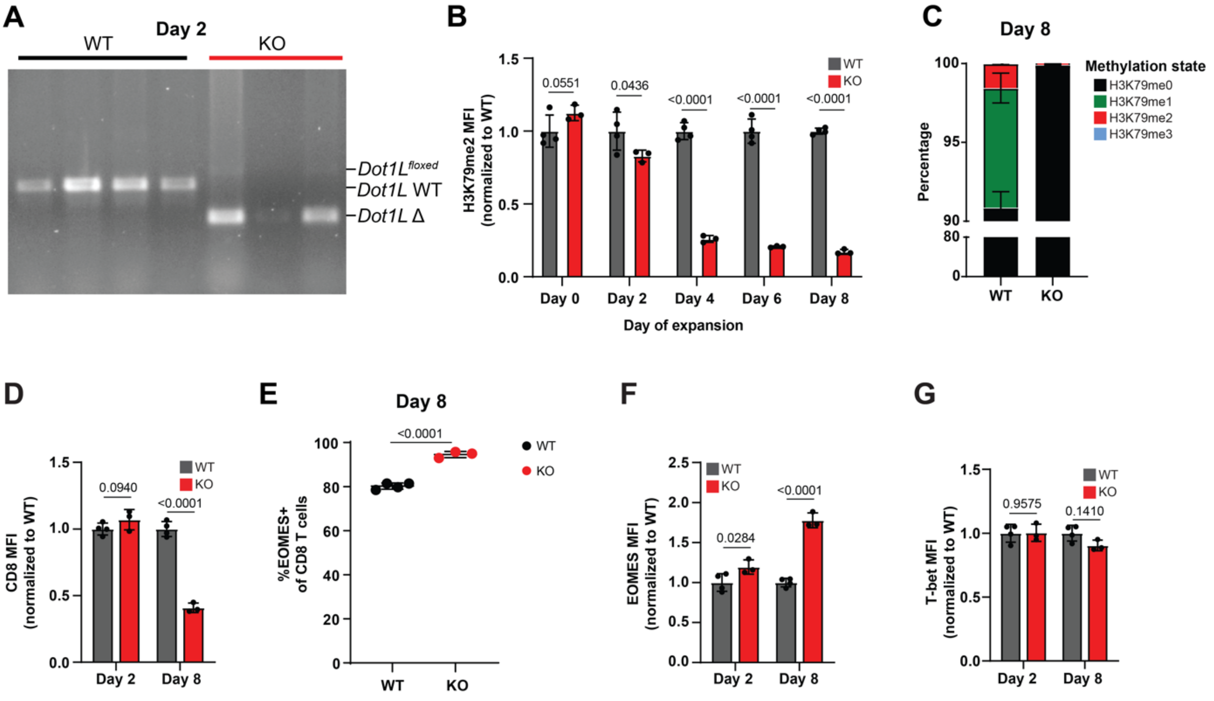


**Figure S1. Confirmation of tamoxifen induced deletion of *Dot1L*. Relates to Figure 1.**
(A) Confirmation of deletion of exon 2 of *Dot1L* for *Dot1L*$^{fl/fl}$ CD8 T cells by PCR, 2 days after 4-OHT treatment *in vitro*. (B) Quantification of H3K79me2 by flow cytometry at during expansion. (C) Confirmation of the loss of H3K79me by mass spectrometry at Day 8 (n=3 biological replicates). (D) Flow cytometry quantification of CD8 at Day 2 and Day 8, normalized to WT. (E) Percentage of EOMES positive CD8 T cells at Day 8. (F-G) Flow cytometry quantification of EOMES (F) and T-bet (G) at Day 2 and Day 8, normalized to WT. Average +/- SD are plotted. n=4 biological replicates for WT and n=3 for KO, unless otherwise indicated. Significance was determined by unpaired t-tests.

### Supplementary Figure S2

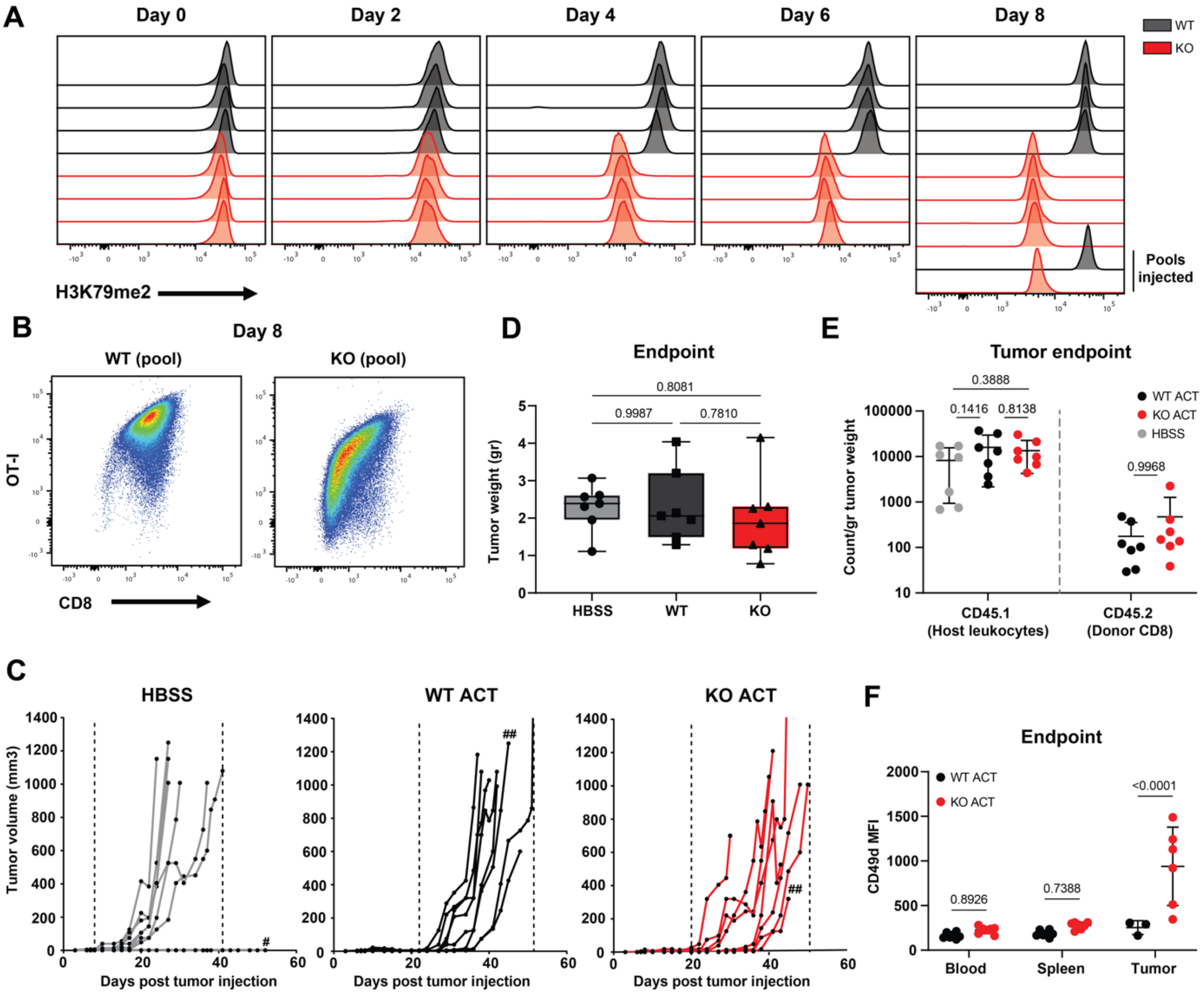


**Figure S2. Tumor control by adoptively transferred *Dot1L* KO CD8 T cells *in vivo*. Relates to Figure 2.** (A) H3K79me2 staining over time (n=4 biological replicates). After 8 days of expansion *in vitro*, CD8 T cells of these mice were pooled for both KO and WT prior to ACT. (B) Flow cytometry dot plots of surface staining for OT-I and CD8 in WT and KO CD8 T cell pools. (C) Tumor outgrowth in individual mice (n=8). Hashtags represents mice excluded from analysis due to lack of tumor growth (#) or death (##). Dotted lines represent time-window of outgrowth. (D) Tumor weight at endpoint. Boxes of the boxplot represent 5-95 percentiles and whiskers represent min-max. Significance was determined by one-way ANOVA and Tukey's multiple comparisons tests. (E) Tumor leukocyte infiltration (count) at endpoint, normalized for tumor weight (average of n=7 +/- SD). (F) Flow cytometry quantification of surface levels of CD49d on donor CD8 T cells in blood, spleen and tumor (average of n=3 (WT) and n=6 (KO) +/- SD; tumor samples with less than 150 events in the CD45.2+ lymphocyte gate were excluded. Significance was determined by 2-way ANOVA and Šídák's multiple comparisons tests, unless otherwise indicated.

#### Supplementary Figure S3

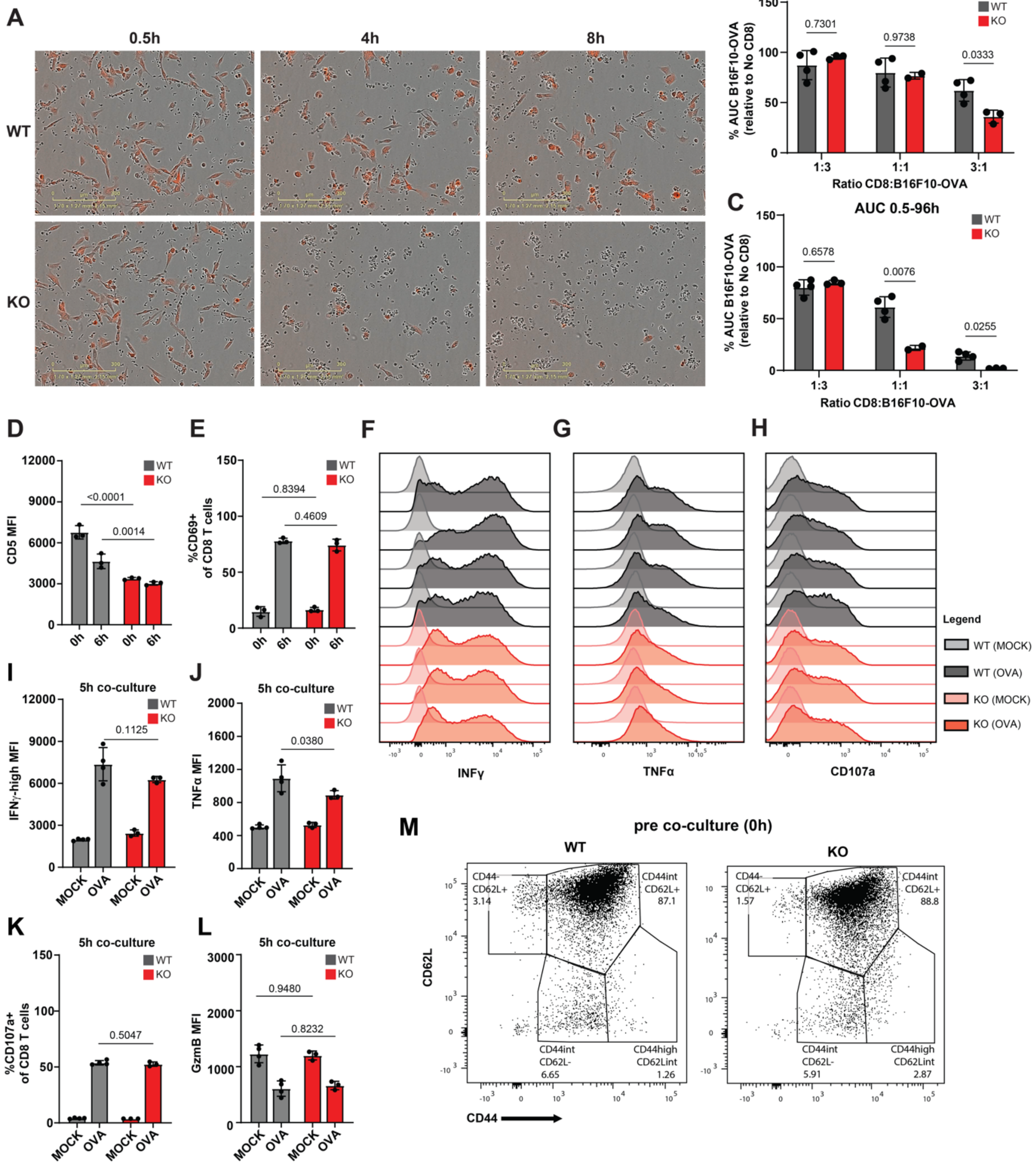

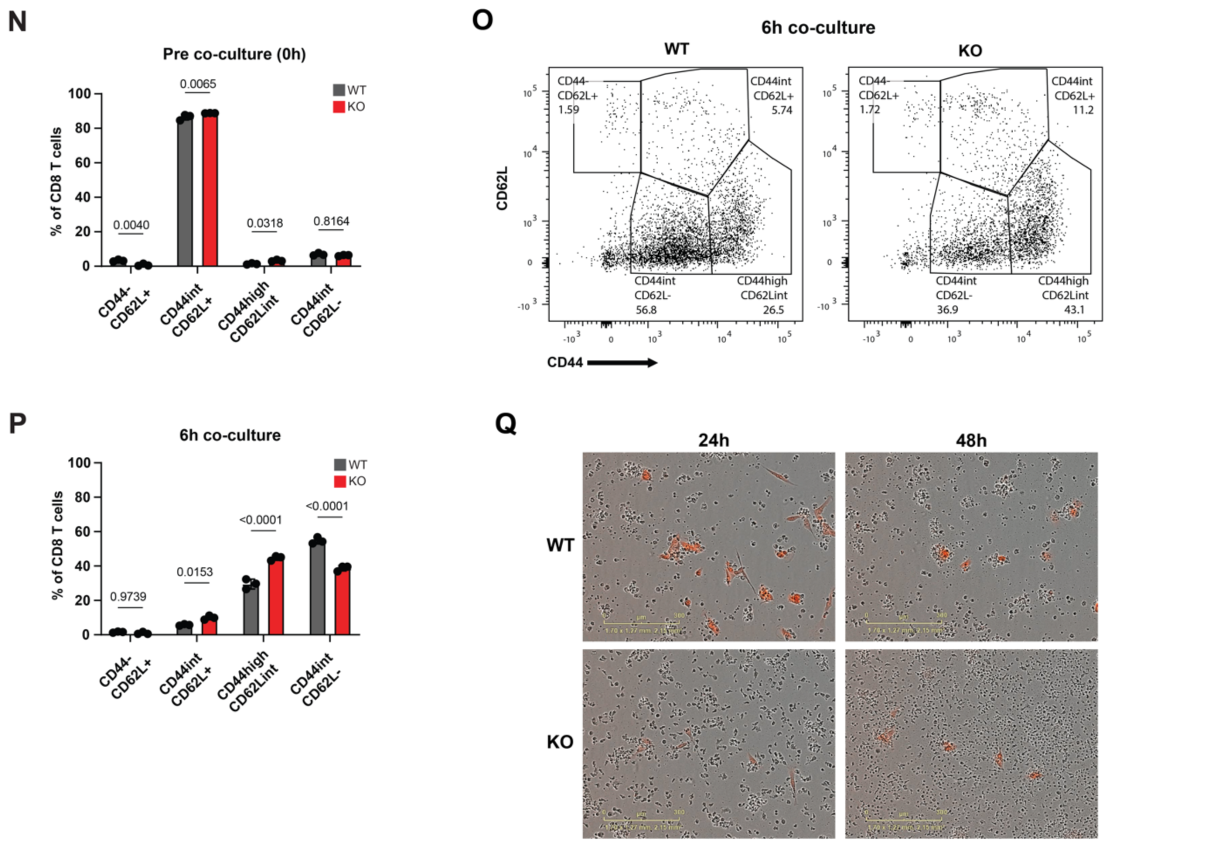


**Figure S3. Deletion of *Dot1L* increases the killing potential of CD8 T cells *in vitro*. Relates to Figure 3.** (A) Live imaging snapshots of B16F10-OVA target cells (red) with WT or KO expanded OT-I CD8 T cells (black dots) over time. Representative cropped images of co-cultures with expanded CD8 T cells (3:1 ratio). (B-C) Relative AUC quantification for 8h and 96h, average of n=3 (KO) and n=4 (WT) +/- SD. (D-E) Quantification of flow cytometry surface level of CD5+ CD8 T cells (D) and %CD69+ (E) pre co-culture and after 6h co-culture (n=3). (F-H) Flow cytometry histograms for effector cytokine production (IFNγ and TNFα) and for degranulation (CD107a) after 5h co-culture with OVA or MOCK B16F10 cells, average +/- SD. (I-L) Quantifications of (F-H) and Granzyme B production after co-culture with OVA or MOCK B16F10 cells for 5h, average of n=3 (KO) and n=4 (WT) +/- SD. (M-P) Representative flow cytometry differentiation dot-plots and quantification of CD44 and CD62L populations for B16F10-OVA co-cultures with WT and KO CD8 T cells, prior to co-culture (M-N) and after 6h co-culture (O-P), average +/- SD (n=3). (Q) Snapshot of live imaging, representative crop of (3:1 ratio) co-culture of B16F10-OVA (red) with expanded WT or KO OT-I CD8 T cells (black dots) at 24h and 48h. Significance was determined by two-way ANOVA and Šídák's multiple comparisons tests.

### Supplementary figure S4

**A**

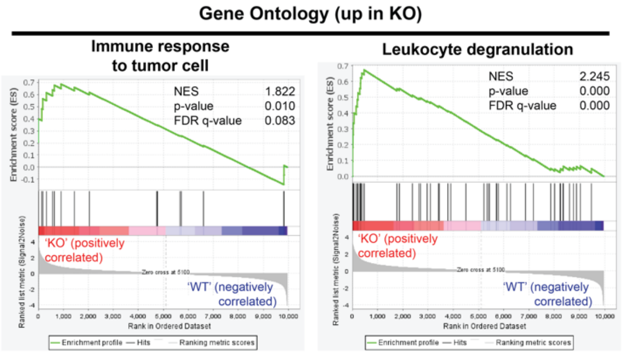


**Figure S4. Genetic ablation of *Dot1L* enriches for upregulation of genes involved in immune response and leukocyte degranulation. Relates to Figure 4.**

(A) GSEA-based GO enrichment for immune response to tumor cell and leukocyte degranulation. Enriched in KO, compared to WT (n=3, biological replicates).

#### Supplementary Figure S5

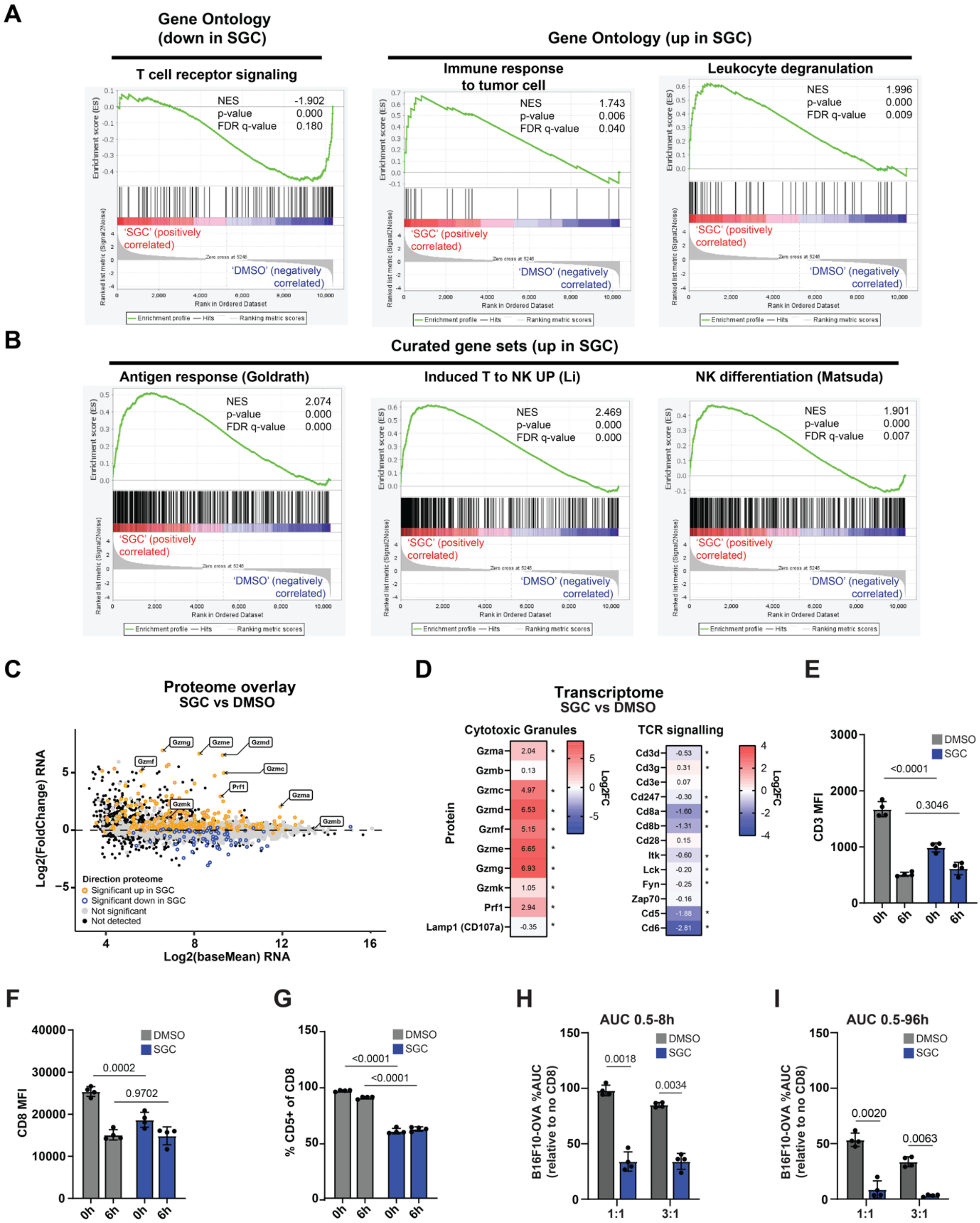


**Figure S5. Loss of catalytic activity of DOT1L drives transcriptomic and proteomic changes. Relates to Figure 5.**

(A) GSEA-based GO enrichment for SGC-0946 (SGC) versus DMSO control (n=4, paired biological replicates). Plotted are T cell receptor signaling, immune response to tumor cells and leukocyte degranulation. (B) GSEA-based curated gene set enrichment. Plotted are antigen experience, induced T to NK and NK differentiation. Normalized enrichment scores (NES), p-values (Padj) and false discovery (FDR) values are displayed. (C) Overlay of the proteome on the transcriptome MA plot. Upregulated (yellow) and downregulated (blue) protein expression are colored accordingly and cytotoxic proteins are highlighted. (D) Heatmaps of selected genes involved in cytotoxic granules and TCR signaling. Asterisk represents adjusted p-value (Padj) significance. (E-G) Flow cytometry of CD8 T cells prior to co culture and after 6h co-culture (n=4, paired biological replicates), showing quantified surface expression of (E) CD3, (F) CD8, and (G) %CD5+. (H-I) Quantification of relative confluency AUC at 8h (H) and 96h (I), relative to no CD8 (n=4, paired biological replicates). (J) Quantified confluency over time of B16F10-OVA treated with SGC-0946 (10 µM), compared to DMSO (n=3 technical replicates). (K) Flow cytometry of CD8 T cells showing percentage of live cells pre co-culture (0h) and after 6h of co-culture. (L-N) Quantification of CD44 and CD62L, and T-bet and EOMES expression states of SGC-0946 and DMSO treated CD8 T cells (flow cytometry), (L-M) prior to co-culture and (N) after 6h co-culture with B16F10-OVA. For E-I, significance was determined by paired two-way ANOVA and Šídák's multiple comparisons tests.

### Supplementary Figure S6

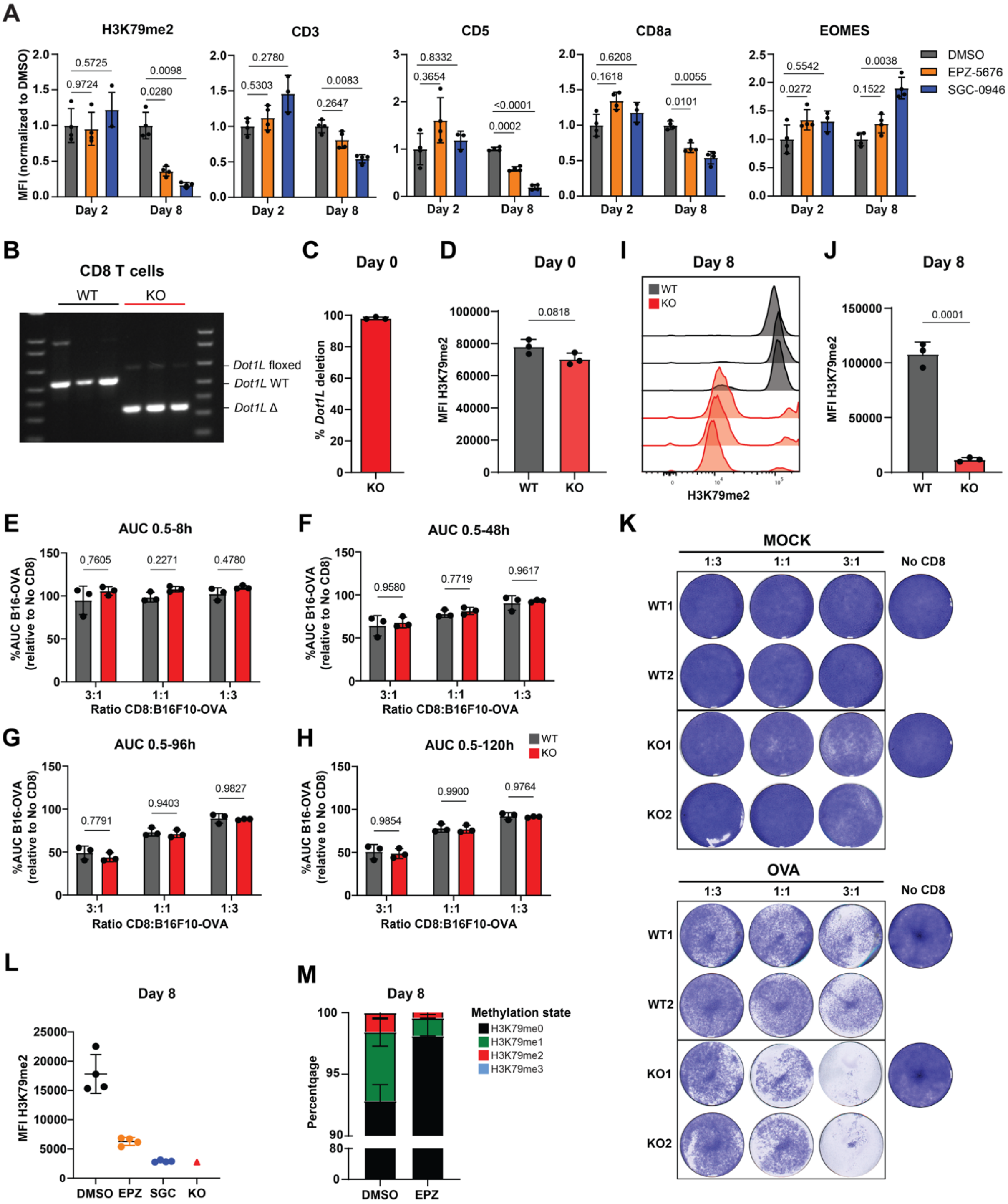

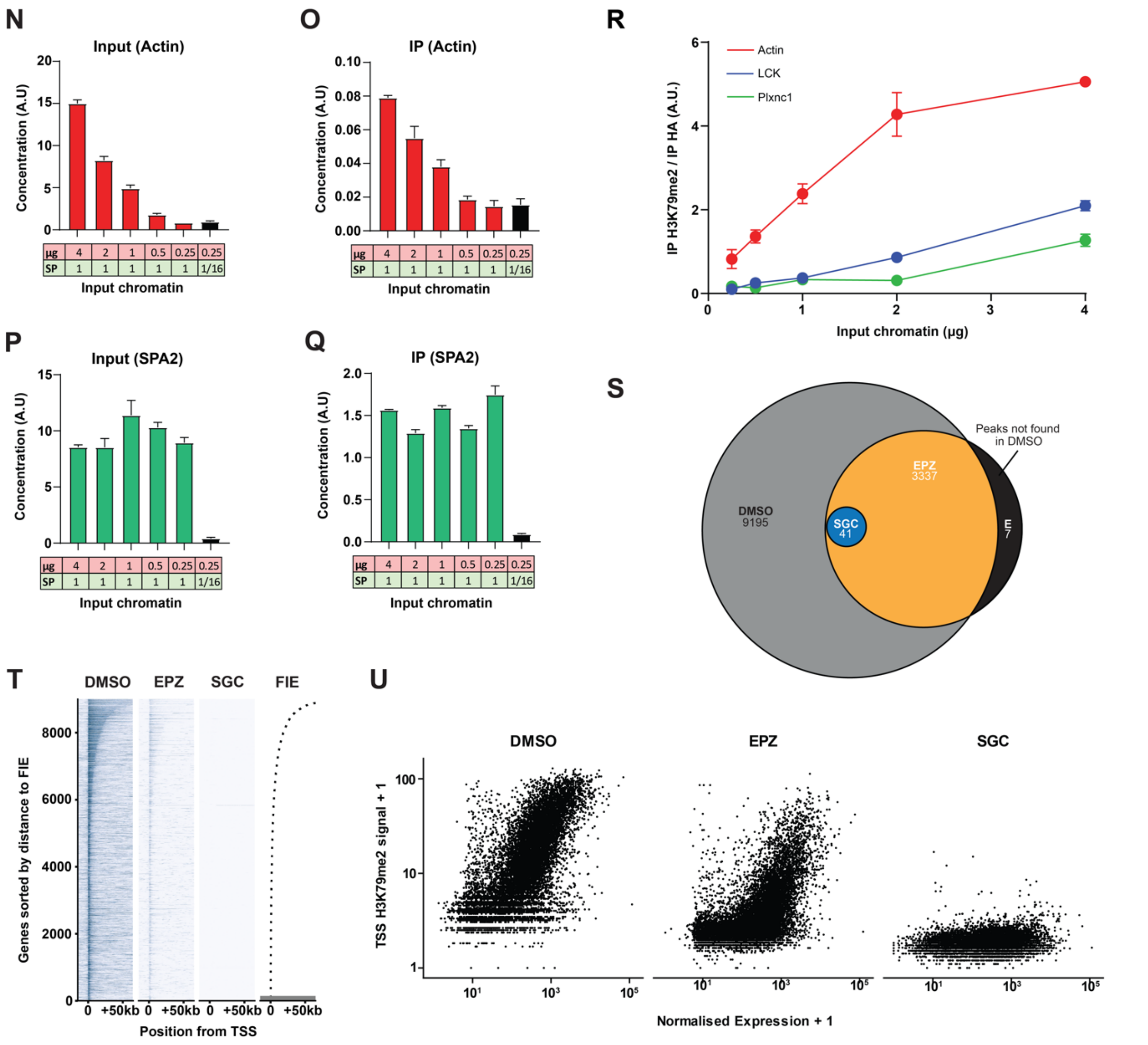


**Figure S6. Loss of H3K79me2 is correlated to increased killing potential, relates to Figure 6.**
(A) Quantification of H3K79me2, CD3, CD5, CD8a and EOMES (flow cytometry) at expansion Day 2 and Day 8. Displayed are the data of CD8 T cells treated with DMSO (0.1 %), EPZ-5676 (10 µM) or SGC-0946 (10 µM) (n=4), normalized to DMSO within each corresponding day. (B) Agarose gel of *Dot1L* PCR on CD8 T cells from Tamoxifen treated mice (N=3). (C) Quantification of (B) to determine deletion efficiency. Average +/- SD. (D) Quantification of figure 6B, average +/- SD (n=3). Significance was determined by unpaired t-test. (E-H) Relative confluency AUC quantifications of live imaging cytotoxicity assay at (E) 8h, (F) 48h, (G) 96h and (H) 120h. Different ratios of CD8 to B16F10-OVA are displayed, average +/- SD (n=3). Significance in relative confluency AUC plots was determined by two-way ANOVA and Šídák's multiple comparisons test, and corresponding P-values were plotted in the graphs. (I) Flow cytometry histogram showing H3K79me2 of CD8 T cells that were derived from tamoxifen treated mice and subsequently expanded for 8 days (n=3). (J) Quantification of I, average +/- SD. Significance was determined by an unpaired t-test. (K) Fixed cytotoxicity assay of expanded KO and WT OT-I CD8 T cells derived from tamoxifen treated mice (n=2). Purple color represents adherent B16F10-OVA cells, lower purple density represents effective killing. (L) Flow cytometry quantification of H3K79me2 at day 8 of treatment with EPZ-5676 (10 µM), SGC-0946 (10 µM) or DMSO (0.1%) (n=4), using Lck-Cre (DOT1L-KO) CD8 T cells (n=1) as a reference. (M) Quantification of H3K79me states by mass spectrometry at Day 8; DMSO and EPZ-5676, average +/- SD (n=4 biological replicates). (N-Q) qPCR analysis of a titration series of variable amounts (µg) of mouse chromatin (red) with a constant yeast chromatin (green) spike-in (SP) using antibodies against HA and H3K79me2, showing data for mouse Actin (N-O) and yeast SPA2 (P-Q) for both input and IP. Plotted are the average of two technical replicates +/- SD. (R) Line plot for H3K79me2 IP, corrected for the average of their respective spike-in SPA2 qPCR values (average of two technical replicates +/- SD). Displayed are three

genes that were selected for H3K79me2 high (Actin), intermediate (LCK) and low (Plxnc1) respectively. (S) Venn diagram of normalized and aggregated (n=2) ChIP peak count quantification for EPZ-5676, SGC-0946 and DMSO treated expanded CD8 T cells. (T) Genomic distribution of H3K79me2 on genes, sorted for H3K79me2 peak intensity near the TSS (0- to +5 kb). (U) Tornado plot H3K79me2 TSS intensity to transcription correlation. Data shown are aggregates of biological replicates (n=2 for H3K79me2, n=4 for expression).
